# Synthetic *in vivo* compartmentalisation improves metabolic flux and modulates the product profile of promiscuous enzymes

**DOI:** 10.1101/2022.11.24.517869

**Authors:** Li Chen Cheah, Lian Liu, Manuel R. Plan, Bingyin Peng, Zeyu Lu, Gerhard Schenk, Claudia E. Vickers, Frank Sainsbury

**Author notes:** Corresponding authors (Frank Sainsbury); (Claudia E. Vickers). Li Chen Cheah: Australian Centre for Disease Preparedness, 5 Portarlington Rd, East Geelong, VIC 3219, Australia. Claudia E. Vickers: Eden Brew, Brisbane QLD 4000, Australia.

## Abstract

Enzyme spatial organisation and compartmentalisation are naturally evolved mechanisms for facilitating multi-step biocatalysis. We explored the synthetic *in vivo* co-encapsulation of two different cargo proteins in yeast using a self-assembling virus-like particle. Co-encapsulation was verified using single particle techniques for both end-to-end fusion of the cargo proteins with the encapsulation anchor at one end, and coexpression of each cargo protein with their individual anchors. The co-encapsulation of a bifunctional geranyl diphosphate/farnesyl diphosphate synthase and a bifunctional linalool/nerolidol synthase delivered nerolidol titres up to 30 times that of an unorganised ‘free’ enzyme control, a remarkable improvement from a single engineering step. Interestingly, striking differences in the ratio of products (linalool and nerolidol) were observed with each spatial organisation approach. This work presents the largest reported titre fold increases from *in vivo* enzyme compartmentalisation and suggests that enzyme spatial organisation could be used to modulate the product profile of promiscuous enzymes.

## INTRODUCTION

In a cell, hundreds of simultaneous metabolic reactions are exquisitely coordinated. A recurring metabolic control strategy in nature is to spatially organise the enzymes that participate in a reaction cascade using metabolons or cellular compartments. Bioinspired strategies such as direct enzyme fusion^1,2^, enzyme organisation with synthetic scaffolds^3–5^, and directing enzymes into membrane-bound organelles^6–8^ are established methods for engineering *in vivo* co-localisation. More recently, heterologous self-assembling protein compartments have been used to spatially organise enzymes in microbial bioproduction pathways^9,10^. Apart from co-localisation functions, compartments are able to sequester the pathway from undesirable interactions that may result in intermediate loss or toxicity^11,12^. The rationale for using completely orthogonal compartments is to further minimise crosstalk with the host cell and allow greater control over the reaction environment^11^.

Artificial metabolic compartments have been constructed by harnessing self-assembling protein cages such as virus-like particles (VLPs)^10,13,14^, bacterial microcompartments^9,15–17^, and encapsulins^18,19^. Despite their size and apparent complexity, these supramolecular structures self-assemble from only a few types of repeating building blocks. Synthetic protein compartments can potentially be engineered for selective metabolite permeability^15,20–22^, enabling applications where accumulation of specific intermediates or the sequestration of toxic metabolites is desired. Protein cage platforms that package heterologous cargo proteins typically have two basic components: (i) the shell/coat protein(s) that form the basic cage structure and (ii) the cargo protein of interest, fused to a targeting peptide, or anchor, that binds to the internal surface of the cage. Another strategy commonly used for cargo packaging is to fuse the cargo protein directly to a lumen-facing terminus of the shell protein^23–26^. However, this approach may interfere with the self-assembly of compartments depending on the size and complexity of the targeted cargo proteins^23,27^.

As many applications of artificial metabolic compartments require the co-encapsulation of two or more enzymes, it is desirable to design co-encapsulation strategies that are efficient, predictable, and tuneable. From the limited body of work on *in vivo*-assembled compartments so far, two broad approaches have been explored for cargo protein co-encapsulation: (i) end-to-end fusion of all the required cargo proteins to a single anchor^13,28^; and (ii) coexpression of each cargo protein fused to its own anchor^10,17,29^. The end-to-end fusion strategy is the simplest to implement; expression is performed exactly as for a single cargo type, using a single promoter for the fusion protein. Theoretically, proteins are encapsulated at a defined 1:1 ratio as genetically encoded. The main drawback of this approach is the ratio of encapsulated proteins cannot be adjusted to suit specific applications. Furthermore, the size of the fusion protein can become very large and unwieldy as more enzymes are added. This may increase the risk of steric clashes that disrupt assembly or decrease loading efficiency. This method also involves tethering both ends of at least one cargo protein, which may not be optimal for enzyme function. The coexpression strategy (*i*.*e*. where each cargo is separately fused to its own anchor) enables more flexibility in adjusting the ratio of encapsulated cargo proteins – for example, by varying gene promoter strength or the binding affinity of the anchors. In addition, this strategy requires each cargo to be tagged only at one end, reducing the risk of disrupting enzyme function.

We have previously developed an *in vivo*-assembling compartment platform for the popular bioproduction chassis, *Saccharomyces cerevisiae*, based on murine polyomavirus (MPyV) VLPs^14^. Each VLP shell is nominally composed of 360 copies of the major capsid protein VP1, which are arranged in 72 pentamers^30^. In the MPyV VLP system, cargo protein encapsulation occurs by tagging the protein of interest with an N-terminal anchor derived from the C-terminus of the minor capsid protein VP2, called VP2C^31^. As a single copy of VP2C can interact with each pentamer of VP1, each VLP can therefore accommodate up to 72 units of the cargo protein. However, since VP2C binding is not essential for VLP assembly, cargo loading *in vivo* is therefore a stochastic process, and capsids can be assembled from a mosaic of cargo-bound and unbound VP1 pentamers. MPyV VLPs are able to encapsulate a diverse range of cargo proteins^14,27^, possibly because the loading density can self-adjust to accommodate larger cargo proteins without preventing assembly. This is in contrast to systems with strict loading stoichiometries (*e*.*g*. where the cargo protein is fused directly to the shell protein), where steric clashes from large cargoes could disrupt self-assembly. The simplicity, modularity, ample loading capacity, and flexibility in accommodating various cargoes make MPyV VLPs an attractive system for metabolic pathway compartmentalisation.

In this work, we directly compare the *in vivo* co-compartmentalisation of two cargo proteins in MPyV VLPs by the end-to-end fusion and coexpression strategies. After verifying that both strategies produced effective cargo co-encapsulation, we applied the strategies to organise two sequential enzymes in a heterologous biosynthesis pathway that produces the sesquiterpene nerolidol. Compartment-forming strains produced much higher product titres compared to the control with unorganised ‘free’ enzymes, demonstrating the potential of MPyV VLPs as an enzyme stabilisation, co-localisation, and metabolic engineering tool. The altered product profiles exhibited by strains with co-encapsulated enzymes also suggest that enzyme spatial organisation may be a novel direction for controlling the specificity of promiscuous enzymes.

## RESULTS AND DISCUSSION

### *In vivo* cargo encapsulation by MPyV VLPs

In the first stage of this work, green and red fluorescent proteins (GFP and mRuby3) were used as model cargo proteins for studying co-encapsulation. We have previously shown that a VP1 mutant that lacks the putative nuclear localisation signal (‘ΔVP1’) exhibits better cargo capture properties and a more diffuse subcellular distribution – indicative of cytosolic localisation – compared to wild-type VP1^14^. The ΔVP1 variant was therefore chosen for this work. Simultaneous expression of ΔVP1 with VP2C-tagged cargo proteins leads to specific *in vivo* packaging of cargo into VLPs. Cargo proteins were either tethered end-to-end (‘Linked’ construct) or individually fused to the VP2C anchor (‘Coexpressed’ construct); schematic diagrams of the expression strategies are shown in Figure 1a. For the Coexpressed construct, different promoters were used for each VP2C-cargo fusion to avoid inadvertent homologous recombination between cassettes^32^. Both strategies produced uniform particles of similar morphology under transmission electron microscopy (TEM; Figure 1b). VLPs arising from the two constructs had very similar particle size distributions, based on nanoparticle tracking analysis (NTA; Figure 1c).

**Figure 1.**
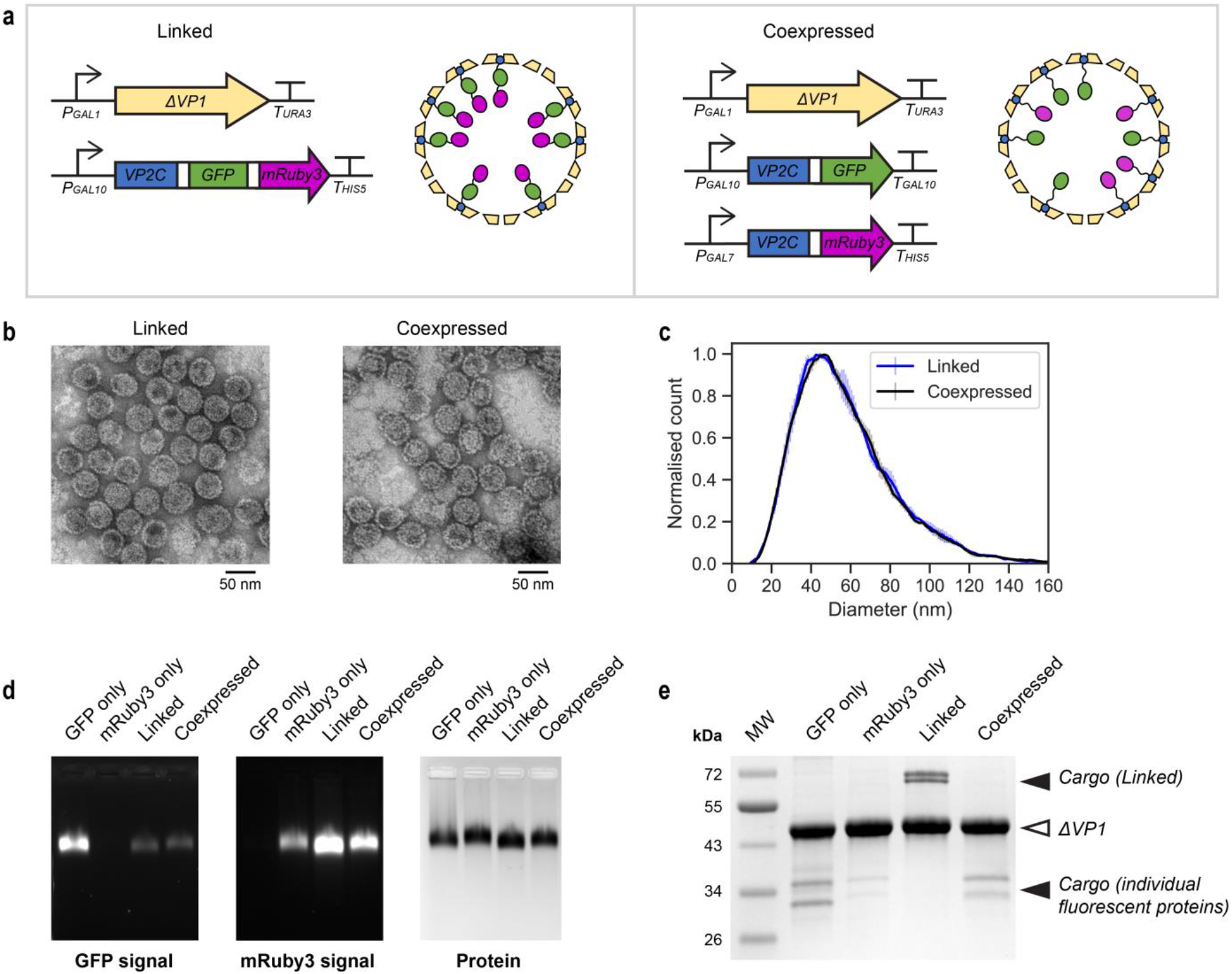
*In vivo* co-encapsulation of red and green fluorescent proteins with the MPyV VLP platform. (a) Schematic of the ‘Linked’ and ‘Coexpressed’ cargo co-encapsulation strategies. Green and red fluorescent proteins (GFP and mRuby3) were used as model cargo. Each domain in the fusion proteins was connected by flexible linkers (shown in white). Representations are not drawn to scale. (b) Negatively stained TEM images of VLPs purified from each co-encapsulation strain. (c) Particle size distributions, measured using NTA. The mean of 3 technical replicates is shown, with error bars indicating +/- 1 standard deviation. (d) Native agarose gel electrophoresis showing GFP and mRuby3 signal in intact particles. Linked and Coexpressed particles were compared with control constructs encapsulating GFP and mRuby3 alone. (e) SDS-PAGE gel stained with Coomassie blue. Arrows show the band positions of ΔVP1 and encapsulated cargo proteins.

Native agarose gel electrophoresis shows the clear signals of both GFP and mRuby3 in intact Linked and Coexpressed VLPs (Figure 1d). Both constructs led to excellent cargo loading, as determined by SDS-PAGE (Figure 1e). Cargo loading estimations by SDS-PAGE densitometry gave 72 units of VP2C-GFP-mRuby3 per Linked VLP (full theoretical capacity) and 61 units of either VP2C-GFP or VP2C-mRuby3 per Coexpressed VLP. Note that the SDS-PAGE shows two cargo bands per construct due to a known spontaneous *in vivo* truncation of VP2C that does not appear to reduce loading of cargo proteins^14^. The Linked construct appeared to encapsulate more mRuby3 than Coexpressed; this is unsurprising given that the theoretical capacity for individual fluorescent proteins is double for Linked VLPs (as two fluorescent proteins share a VP2C anchor in the Linked configuration). The higher mRuby3 signal on native agarose gel electrophoresis from Coexpressed VLPs compared to the mRuby3-only control VLPs may be explained by the use of a stronger promoter (P_GAL7_ *versus* P_GAL10_)^32^.

Curiously, we consistently observed much better loading of GFP-containing cargo proteins compared to cargo without GFP (*i*.*e*. VP2C-mRuby3). The disparity is especially evident in the control constructs that encapsulate either VP2C-GFP or VP2C-mRuby3 alone, which were expressed using the same promoter (Figure 1e). *In vivo* cargo loading is not well understood and non-specific charge interactions between GFP and the positively charged internal surface of the MPyV shell may affect cargo loading; the GFP variant has a net charge of -7 (34 negatively charged residues and 27 positively charged residues), while mRuby3 only has a net charge of -4 (33 negatively charged residues and 29 positively charged residues). Another possibility is that the weak dimerisation tendency of GFP^33^ may have influenced the assembly process.

### Effective fluorescent protein co-encapsulation by both Linked and Coexpressed approaches

Ensemble measurement techniques cannot measure the degree of co-encapsulation or the spatial arrangement of the two cargo proteins within individual VLP. We thus examined if co-encapsulated fluorescent proteins are in close proximity using Förster resonance energy transfer (FRET). For fluorescent proteins, efficient energy transfer only occurs if the donor and acceptor are within approximately 10 nm^34^. Our previous work on *in vitro*-assembled MPyV VLPs, using eGFP as the donor and mRuby3 as the acceptor^31^, demonstrated the occurrence of FRET between cargo proteins spaced ∼7 nm apart. Accordingly, the emission spectra of both constructs (at 450 nm excitation) show a clear FRET acceptor peak in the mRuby3 emission range (Figure 2a). Control VLPs with either GFP or mRuby3 alone did not show a significant FRET peak with the same settings (Supporting Information, Figure S1). Moreover, mixing the control VLPs did not increase the signal in the acceptor range, indicating that co-encapsulation is required for significant energy transfer to occur (Figure S1). Coexpressed VLPs exhibited slightly lower FRET than Linked VLPs, presumably due to an uneven distribution of GFP and mRuby3 between and within VLPs.

**Figure 2.**
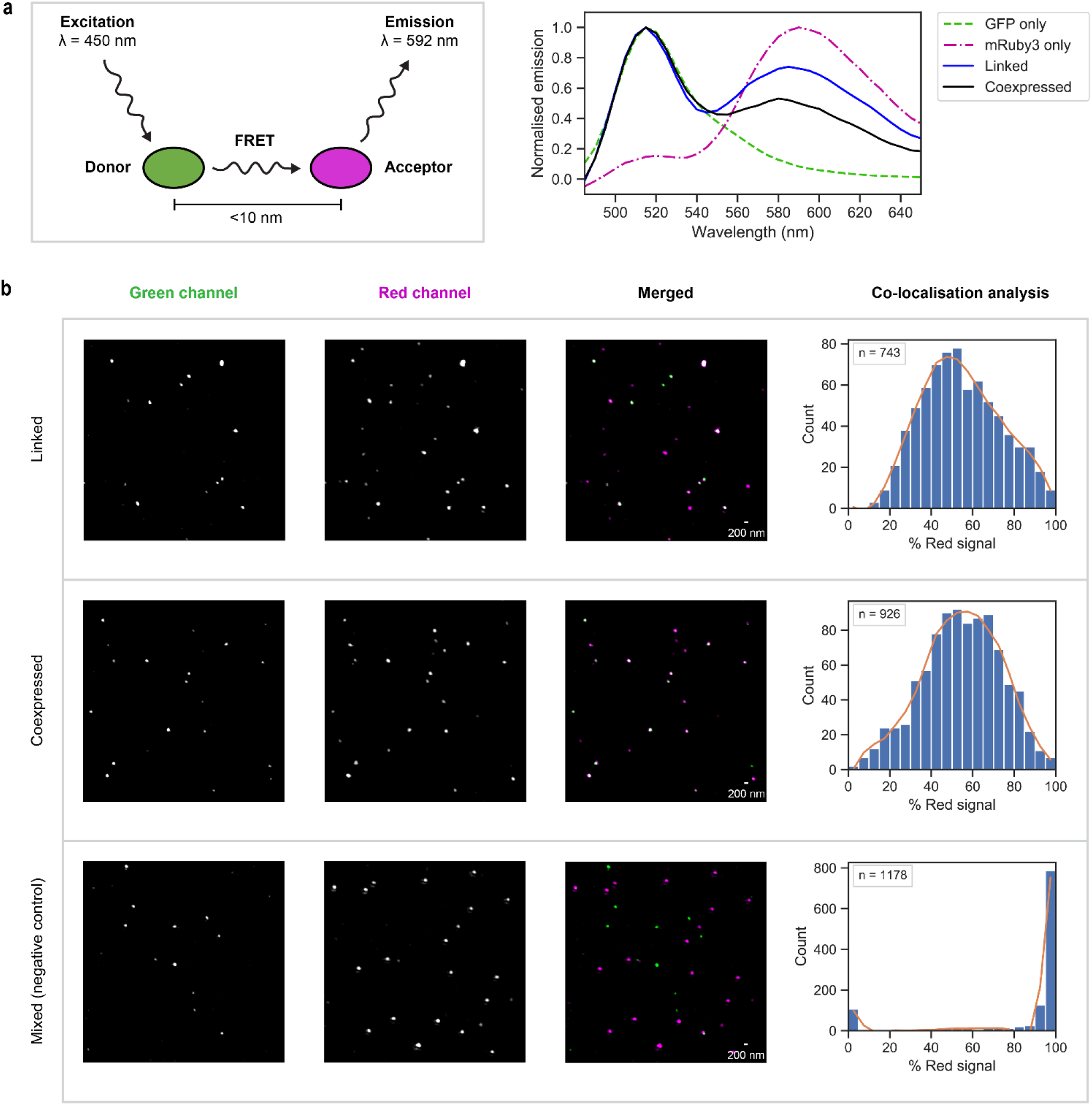
Assessing the degree of cargo co-encapsulation with single-particle techniques. (a) Förster resonance energy transfer (FRET) assay, with GFP as the donor and mRuby3 as the acceptor. Efficient energy transfer only occurs if the fluorophores are within <10 nm of each other. The assay was performed at 450 nm excitation and the emission maximum of mRuby3 is 592 nm. The emission spectra (mean of 3 replicates) have been baseline-subtracted and normalised to each of their respective maximum peaks. Both the Linked and Coexpressed spectra show clear GFP-mRuby3 FRET. Direct excitation of the mRuby3-only control only produces minimal signal; refer to Figure S1 for examples of raw emission spectra. The mean of 3 technical replicates is shown. (b) Imaging of individual VLPs by superresolution structured illumination microscopy (SR-SIM). In the Merged images, the GFP signal is coloured green and the mRuby3 in magenta, which overlap to produce white. For clarity, both the brightness and contrast have been increased; see Figure S2 for an example of an uncropped image visualised with the settings used for data analysis. Quantitative co-localisation analysis reveals similar, high levels of signal co-localisation for Linked and Coexpressed; in contrast, the negative control does not exhibit substantial signal co-localisation. The histograms show the proportion of red signal (from mRuby3) to the total signal (Green + Red) from each individual particle. Histograms were plotted with bin width = 5, and the number of particles in the dataset (n) is shown as an inset.

We assessed the co-encapsulation ratio of GFP and mRuby3 in individual VLPs using super-resolution structured illumination microscopy (SR-SIM), a single-particle imaging technique (Figure 2b). The multicolour imaging capabilities and ability to use standard fluorescent proteins make SR-SIM well-suited for this purpose. A similar SR-SIM protocol was previously used to examine MPyV VLPs assembled *in vitro*^31^. With sufficient dilution, VLPs can be dispersed and adhere evenly onto a glass coverslip. VLPs immobilised at the buffer-glass interface were imaged using 3D SR-SIM. The red fluorescence signal of each particle was divided by the total signal intensity (green + red) of that particle, providing a measure of the mRuby3:GFP ratio.

Both co-encapsulation strategies produced very similar, approximately normal distributions of the mRuby3:GFP ratio (represented as % red signal; Figure 2b), indicating effective fluorescent protein co-encapsulation. Remarkably, the stronger promoter used for VP2C-mRuby3 expression in the Coexpressed strain was able to compensate for its lower encapsulation alone, when compared to VP2C-GFP (Figure 1e), leading to a degree of co-encapsulation comparable to direct protein fusion. This was not observed in the negative control sample (mixed GFP-only and mRuby3-only VLPs), confirming that co-localisation was within particles and not due to low-order aggregates of VLPs. In contrast to previous findings on *in vitro*-assembled MPyV VLPs, which found an apparent bimodal distribution of the mRuby3 signal^31^, we did not observe partitioning of each cargo type into separate particles. This indicates that there is efficient mixing and co-assembly of VLP components *in vivo*.

### Enzyme co-encapsulation greatly improves the productivity of a metabolic pathway

We next sought to assess the utility of both co-encapsulation approaches for organising enzymes that catalyse successive reaction steps in a model metabolic pathway, specifically the biosynthesis of the sesquiterpene nerolidol. Nerolidol is an industrially relevant compound used as a perfume and flavour additive^35^. Extraction from natural sources or total chemical synthesis is costly and has poor yields^35^, making nerolidol a suitable target for bioproduction using microbial cell factories. *S. cerevisiae* contains a native mevalonate pathway, which produces precursors for the synthesis of the cell membrane component ergosterol (Figure 3a). The isoprenoid intermediates geranyl diphosphate (GPP, C10) and farnesyl diphosphate (FPP, C15) are produced by successive additions of isopentenyl diphosphate (IPP, C5) to dimethylallyl diphosphate (DMAPP, C5). Both addition steps are catalysed by yeast FPP synthase (FPPS, also called ERG20). A proportion of FPP is further condensed with another IPP unit to make geranylgeranyl diphosphate (GGPP, C20), catalysed by BTS1. In the ergosterol pathway, FPP is converted into squalene by squalene synthase (SQS). However, FPP can be redirected towards the production of sesquiterpenes by the introduction of a heterologous sesquiterpene synthase such as *Actinidia chinensis* (golden kiwifruit) nerolidol synthase^36^ (NES; Figure 3a). This variant of NES is promiscuous and can also convert GPP into linalool as a minor product^36^. Since terpene synthases are generally known to be slow enzymes^37^, NES is suspected to be one of the rate-limiting enzymes in nerolidol bioproduction. We hypothesised that the co-encapsulation of FPPS with NES could potentially facilitate more efficient redirection of metabolic flux through to NES as well as reduce competition with the native enzymes SQS and BTS1 for the FPP pool.

**Figure 3.**
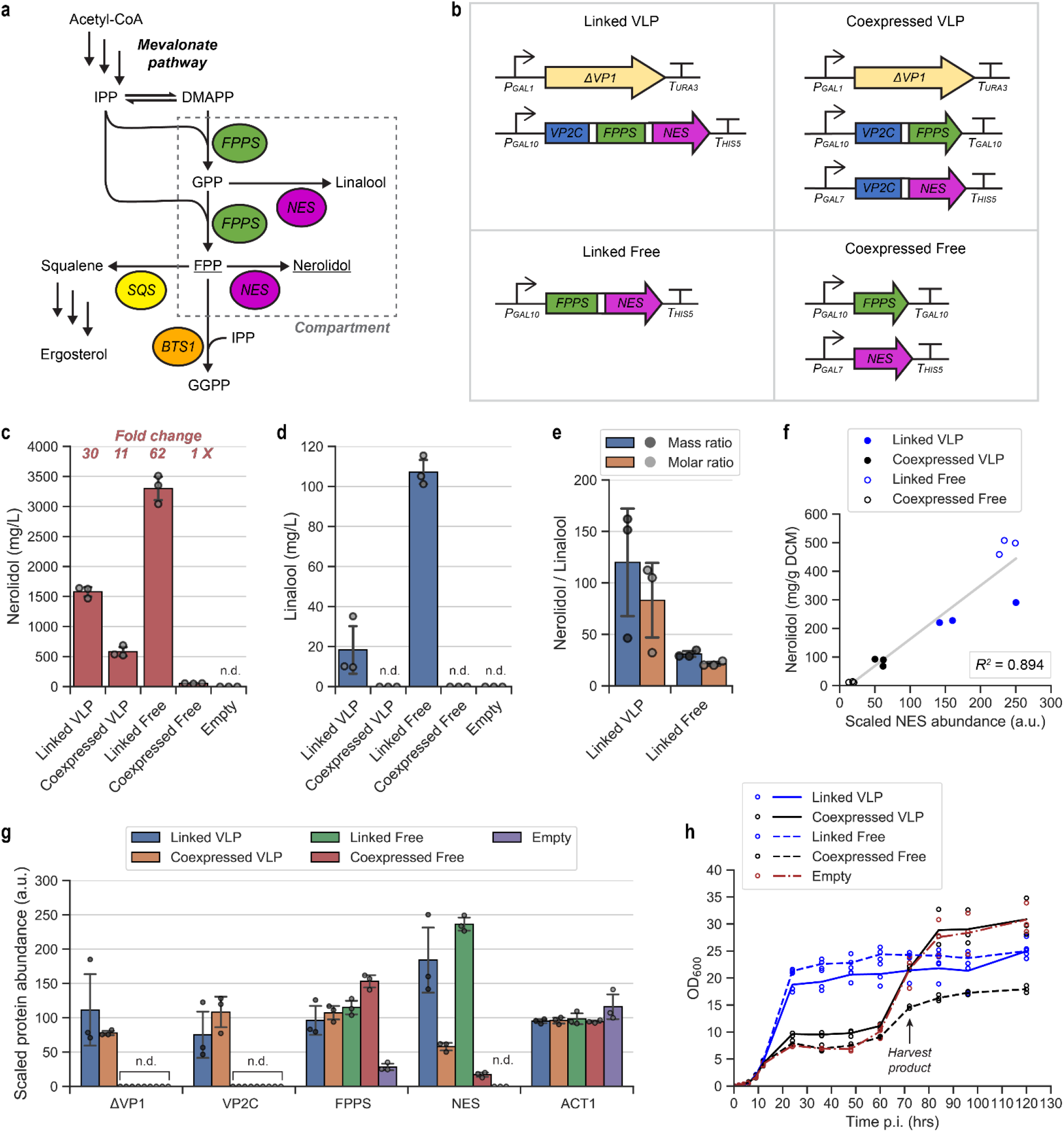
Co-encapsulation of farnesyl diphosphate synthase (FPPS) and nerolidol synthase (NES). (a) The yeast mevalonate pathway produces isopentenyl diphosphate (IPP) and dimethylallyl diphosphate (DMAPP), the building blocks of all isoprenoids. Farnesyl diphosphate (FPP) is formed by two successive additions of IPP to DMAPP by farnesyl diphosphate synthase (FPPS) via the intermediate geranyl diphosphate (GPP). *S. cerevisiae* natively produces FPP as the precursor of squalene (produced by squalene synthase, SQS), an intermediate in ergosterol biosynthesis. FPP is also converted into geranylgeranyl diphosphate (GGPP) by the native enzyme geranylgeranyl diphosphate synthase (BTS1). The heterologous enzyme nerolidol synthase (NES) converts FPP into nerolidol and GPP into linalool. The major products of FPPS and NES (FPP and nerolidol respectively) are underlined. In this study, we examined the compartmentalisation of FPPS and NES (dotted grey box) as a strategy to organise the nerolidol production pathway and reduce competition with SQS and BTS1. (b) Four constructs for expressing FPPS and NES were generated: two for VLP co-encapsulation (‘Linked VLP’ and ‘Coexpressed VLP’) and two non-VLP forming, ‘free enzyme’ controls (‘Linked Free’ and ‘Coexpressed Free’). ‘Empty’ is the base strain transformed with an empty vector. (c) Nerolidol titres at 72 h, in mg per L cell culture. ‘Fold change’ indicates the relative titre compared to the Coexpressed Free strain. Means and individual data points are shown, with error bars +/- 1 STD; ‘n.d.’ = not detected. (d) Linalool titres at 72 h, in mg per L cell culture. Means and individual data points are shown, with error bars +/- 1 STD; ‘n.d.’ = not detected. (e) Mass and molar ratios of the NES products nerolidol and linalool. Means and individual data points are shown, with error bars +/- 1 STD. Product ratios of Coexpressed VLP and Coexpressed Free cannot be reliably calculated because linalool was below the limit of detection. (f) Relationship between nerolidol production (normalised to dry cell mass, mg/g DCM) and relative NES expression. Fitting a linear regression model using the least squares method produces *R*^*2*^ = 0.894. (g) Relative *in vivo* levels of key proteins in the engineered nerolidol pathway at 72 h p.i., as determined by LC-MS/MS. To facilitate comparisons, the abundance of each protein has been scaled so that the average is 100 a.u. across strains. ACT1 (actin) is included as a housekeeping reference protein. The unusually high gene expression in one of the three Linked VLP biological replicates was confirmed to be the result of multiple copy genomic integration (see Figure S8). Means and individual data points are shown, with error bars +/- 1 STD; ‘n.d.’ = not detected. (h) Cell density (measured using OD_600_) over the course of fermentation. Time points are hours post-inoculation (p.i.). Cultures were harvested at 72 h p.i. for metabolite analysis, as indicated by the arrow. Means and individual data points are shown.

To assess the effect of FPPS and NES co-encapsulation, two VLP-expressing constructs were generated (‘Linked VLP’ and ‘Coexpressed VLP’) (Figure 3b). To ensure detectable nerolidol production, the cassettes were transformed into a base strain with a highly upregulated mevalonate pathway (o57BR^38^). The native copy of the yeast FPPS gene was retained in all strains. Co-encapsulation of FPPS and NES increased nerolidol production by ∼30-fold (Linked VLP) and ∼11-fold (Coexpressed VLP) relative to a strain expressing unorganised, free enzymes (Coexpressed Free; Figure 3c), indicating greatly enhanced metabolic flux through the NES pathway. The combination of flux push (by highly overexpressing the mevalonate pathway in the base strain^38^) and possible flux pull from FPPS and NES co-encapsulation resulted in exceptionally high product titres, even without optimising expression levels or fermentation conditions. To our knowledge, this is the first demonstration of metabolic enzyme co-encapsulation in a self-assembling synthetic yeast compartment, and represents the largest titre fold increases reported for engineered *in vivo* compartmentalisation. The nerolidol titres were 1.58 and 0.58 g/L culture for Linked VLP and Coexpressed VLP respectively, which are exceptionally high levels using non-optimised shake-flask conditions. FPPS-NES fusion alone (Linked Free), however, delivered a ∼62-fold increase at 3.3 g/L. We have previously reported on the unusually high nerolidol production of this^39^, which is caused by the apparent stabilisation of NES by FPPS-NES fusion that we also observe here (Figure 3g). We thus hypothesise that the substantial improvements in nerolidol titre in the two VLP-forming strains is due to the stabilisation of NES by compartmentalisation.

We performed whole-cell proteomics on samples collected at 72 h p.i. to examine if the product titres could be attributed to differences in intracellular FPPS and NES levels. As with our previous work^39^, nerolidol production roughly correlates with NES expression (Figure 3f) – NES expression was highest in the Linked Free construct, followed by the Linked VLP, Coexpressed VLP and Coexpressed Free constructs (Figure 3g). There was a >3-fold higher level of NES in Coexpressed VLP compared to Coexpressed Free, despite being fused to the VP2C anchor that is known to destabilise cargo proteins prior to VLP assembly^14^. This observation supports our hypothesis that NES has been stabilised by VLP encapsulation. A truncated version of ΔVP1^40,41^ which forms pentamers and binds VP2C^42^, but does not self-assemble into VLPs, fails to stabilise cargo proteins *in vivo* (Figure S9); this demonstrates that cargo proteins are stabilised by compartmentalisation resulting from VLP assembly, instead of VP2C-ΔVP1 binding.

Proteomics also revealed that one biological replicate of Linked VLP had anomalously elevated expression of the engineered nerolidol pathway (Figure 3g, Figure S6). Quantitative PCR on yeast genomic DNA confirmed that this replicate contained two integrated copies of the entire nerolidol expression cassette (Figure S8). Interestingly, this replicate produced a similar amount of nerolidol but much more linalool (lower nerolidol:linalool ratio) compared to the two other Linked VLP replicates (Figure 3c), with no effect on growth. Although we are not able to explain this finding, it suggested that the relative stoichiometry of these promiscuous enzymes could affect the product profile of the metabolites they produce. This is an understudied phenomenon in synthetic biology, yet one that could have an important impact on the efficiency of metabolite recovery and productivity of cellular biocatalysis.

Consistent with our previous work^39^, the Coexpressed Free and empty vector controls showed impaired growth in the early phase of fermentation (Figure 3h), possibly from the toxic accumulation of isoprenoid pathway intermediates^43,44^. FPPS and NES co-localisation by protein fusion was able to partly alleviate this toxicity, resulting in relatively normal growth of the Linked strains. Coexpressed VLP exhibited an unusual growth profile where growth was initially slow but recovers after 72 h, suggestive of insufficient NES levels to ‘mop up’ excess FPP at the early stages of fermentation followed by an easing of the metabolic flux bottleneck later in the fermentation.

Squalene and ergosterol levels were not significantly changed by VLP expression (Figure S5), despite the sequestration of FPPS away from SQS in the pathway design. We presume that this is because the native copy of FPPS (which was retained in all strains) already supplies sufficient FPP for cell growth and maintenance, meaning the strains would not be drawing from the extra FPP produced by the engineered pathway. Furthermore, the ergosterol pathway is known to be tightly regulated in yeast to prevent the accumulation of toxic sterol intermediates^45,46^. We also detected the common isoprenoid pathway side products farnesol (19.2–48.1 mg/L) and geranylgeraniol (0.71–2.98 mg/L) in all cultures (Figure S5). Given their relatively low amounts compared to nerolidol, nonspecific conversion to prenyl alcohols was thus not considered a significant source of metabolite loss.

### Co-encapsulation of promiscuous metabolic enzymes alters the product profile

The ability of NES to use GPP as an alternative substrate, producing linalool, creates competition with FPPS at the GPP node (Figure 3a). Linalool production was not detected for the two constructs with unfused enzymes (Coexpressed VLP and Coexpressed Free). However, the FPPS-NES fusion constructs Linked VLP and Linked Free produced a substantial amount and there were striking differences in the ratio of nerolidol and linalool. Encapsulation of the fusion (*i*.*e*. Linked VLP) increased the selectivity for nerolidol production by almost four-fold relative to Linked Free (Figure 3e). The capability to tune the product selectivity of a promiscuous enzyme using a localisation or confinement strategy represents a novel approach to addressing a key bioproduction challenge, as the accumulation of a metabolite at high purity (*e*.*g*. nerolidol) would reduce the requirement for further downstream purification.

To further investigate the impact of co-encapsulation on the product profile of NES, we generated a set of strains where FPPS was replaced with a GPP-overproducing FPPS variant (F96W-N127W double mutant^47^) in the galactose-inducible cassettes. This variant of FPPS has a much lower affinity for GPP compared to the wild-type enzyme (K_M_^GPP^ = 27.6 μM *vs* 0.43 μM, respectively)^47^, which would enhance NES competitiveness for the GPP pool and is expected to increase the linalool titre. Comparisons of strains expressing the GPP-overproducing FPPS variant (Figure 4b) with the wtFPPS-expressing set (Figure 3d) confirms that this was indeed the case, but with nerolidol still being produced at an order of magnitude higher titre (Figure 4a). This is likely due to the strong preference of NES for FPP compared to GPP, with k_cat_/K_M_ values of 300 s^-1^ mM^-1^ for FPP and 69 s^-1^ mM^-1^ for GPP^36^. Additionally, the higher volatility of the C10 product linalool could have led to greater loss by vaporisation compared to the C15 product nerolidol. It is not known how much FPP is still produced by the F96W-N127W mutant; in any case, nerolidol production would also be supported by FPP production by native FPPS (which was retained in the base strain). Interestingly, Linked VLP produced the highest nerolidol titre among the strains using this FPPS variant, at 2.2 g/L (Figure 4a). No obvious growth impairments were exhibited by any of the strains expressing FPPS(F96W-N127W) and NES (Figure 4c). In the event of a metabolic flux bottleneck, strains overexpressing wtFPPS are expected to accumulate higher intracellular concentrations of FPP than strains overexpressing the GPP-overproducing variant. We thus inferred that – at least in this strain background – FPP accumulation poses a higher metabolic burden than GPP accumulation.

**Figure 4.**
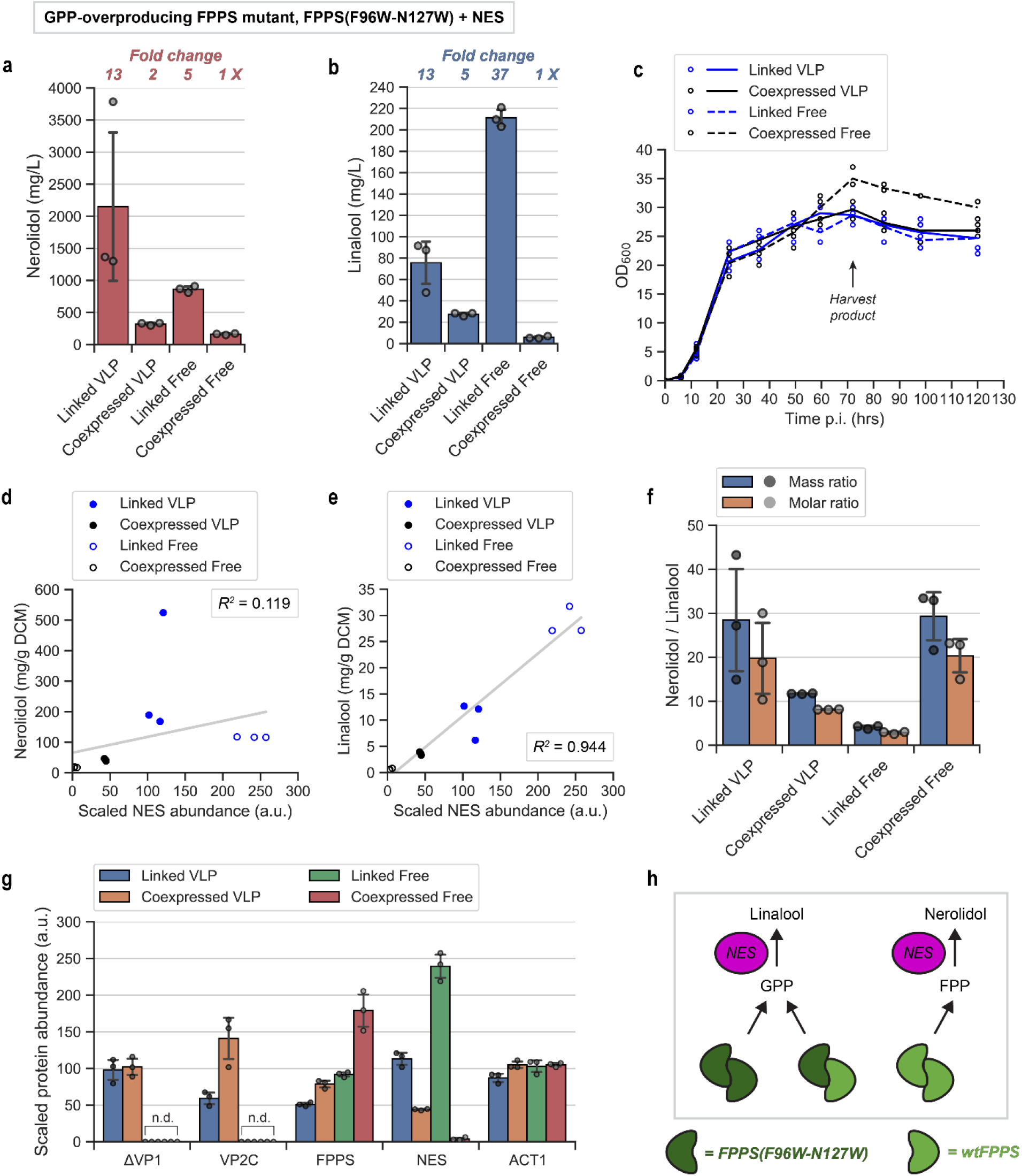
Co-encapsulation of a GPP-overproducing FPPS variant (F96W-N127W) and NES. (a) Nerolidol titres at 72 h, in mg per L cell culture. ‘Fold change’ indicates the relative titre compared to the Coexpressed Free strain. Means and individual data points are shown, with error bars +/- 1 STD; ‘n.d.’ = not detected. (b) Linalool titres at 72 h, in mg per L cell culture. ‘Fold change’ indicates the relative titre compared to the Coexpressed Free strain. Means and individual data points are shown, with error bars +/- 1 STD; ‘n.d.’ = not detected. (c) Cell density (OD_600_) over the course of fermentation. Cultures are harvested at 72 h p.i. for metabolite analysis, as indicated by the arrow. Means and individual data points are shown. (d) Relationship between nerolidol production (normalised to dry cell mass, mg/g DCM) and relative NES expression. Fitting a linear regression model using the least squares method produces *R*^*2*^ = 0.119. (e) Relationship between linalool production (normalised to dry cell mass, mg/g DCM) and relative NES expression. Fitting a linear regression model using the least squares method produces *R*^*2*^ = 0.944. (f) Mass and molar ratios of the NES products nerolidol and linalool. Means and individual data points are shown, with error bars +/- 1 STD. (g) Relative *in vivo* levels of key proteins at 72 h p.i., as determined by LC-MS/MS. The abundances are scaled so that the average across all strains is 100 a.u. to facilitate comparisons. ‘FPPS’ abundance includes both the overexpressed F96W-N127W mutant and native FPPS, which are indistinguishable due to their high sequence identity. ACT1 (actin) is included as a housekeeping reference protein. Means and individual data points are shown, with error bars +/- 1 STD; ‘n.d.’ = not detected. (h) Heterodimers of FPPS(F96W-N127W) and wtFPPS produce predominantly GPP, and so does homodimers of FPPS(F96W-N127W). This means that FPP, and in turn, nerolidol production is mainly supported by homodimers of native FPPS. Enzyme spatial organisation could potentially alter the proportion of FPPS heterodimers and influence the ratio of metabolite products, for example by sequestering FPPS(F96W-N127W) subunits away from the native FPPS pool.

Metabolite production by the FPPS(F96W-N127W) strains resembles the trends observed for the wtFPPS strains. In this case, linalool production increases proportionally with NES expression levels (Figure 4e). Whole-cell proteomic analysis revealed a similar trend in protein expression to wtFPPS-overexpressing strains (Figure 4g). As before, NES expression was greatly enhanced by VLP encapsulation and enzyme fusion. Concomitantly, the highest linalool titres were obtained from the Linked Free construct, followed by Linked VLP and Coexpressed VLP. Relative NES expression of the Coexpressed Free configuration was even lower than with wtFPPS, which could indicate differences in *in vivo* stability between the two FPPS variants. The production of both nerolidol and linalool was enhanced by encapsulation and stabilisation of the coexpressed enzymes.

There is a curious exception to the correlation between NES stabilisation and terpene production. Using the mutant FPPS(F96W-N127W), Linked VLP increased the selectivity for nerolidol production by almost seven-fold compared to ‘Linked Free’ (Figure 4f). Unlike before with wtFPPS overexpression, this was not just due to the decrease in linalool production relative to ‘Linked Free’, but was also a result of higher nerolidol production. A potential side effect of enzyme spatial organisation is the sequestration of mutant FPPS from native FPPS. Yeast FPPS is predicted to be homodimeric, based on homology modelling and comparisons with the structure of avian FPPS^47^. The FPPS(F96W-N127W) mutant was designed to be dominant negative, where heterodimerization with a wtFPPS subunit causes the complex to behave like the mutant version and overproduce GPP^47^ (Figure 4h). Therefore, GPP would be produced by a combination of FPPS(F96W-N127W) homodimers and the heterodimer with native FPPS. This would support high linalool production in the case of Linked Free. Crucially, the encapsulation of FPPS(F96W-N127W) by VLPs may, to some extent, inhibit the formation of heterodimers with native FPPS, thus promoting FPP production by native FPPS and, in turn, nerolidol production.

### Enzyme spatial organisation could influence substrate accessibility

The spatial organisation approaches that we explored produced a clearer effect on competition at the GPP node (FPPS *vs* NES) compared to competition at the FPP node (NES *vs* SQS/BTS1). Although the original motivation for FPPS and NES compartmentalisation was to reduce competition from the native enzymes SQS and BTS1 for the intermediate FPP, any discernible effects were masked by the tight regulation of the ergosterol pathway, and the fact that only a very small proportion of the FPP pool is being utilised by SQS and BTS1. In our strains, nerolidol and linalool production were also strongly tied to the level of NES (Figure 3g, Figure 4g), which is the rate-limiting enzyme. VLP encapsulation and/or enzyme fusion led to *in vivo* stabilisation of NES, which in turn alleviated growth impairments caused by the toxic accumulation of isoprenoid intermediates (Figure 3h) and, ultimately, increased product titres.

Overall, configurations where FPPS and NES are translationally fused (Linked VLP and Linked Free) resulted in the highest linalool titres. Intriguingly, VLP compartmentalisation produced opposing effects on linalool production depending on the co-encapsulation strategy. The Linked VLP configuration more than halved linalool production compared to Linked Free, while the Coexpressed VLP configuration increased linalool production by almost five-fold compared to the Coexpressed Free control (for FPPS(F96W-N127W)). The influence of compartmentalisation on competition for the intermediates GPP (10C) and FPP (15C) parallels a recent yeast study, where relocating the enzymes in a recursive alcohol elongation pathway from the cytoplasm to the mitochondrion favoured accumulation of the longer chain product isopentanol (5C) over isobutanol (4C)^8^. These observations suggest the utility of enzyme spatial organisation as a tool to modulate the accessibility of an intermediate metabolite in order to drive a preferred product profile.

There are multiple, often overlapping explanations for how substrate accessibility can be altered by enzyme co-localisation and spatial organisation. Intuitively, bringing successive acting enzymes in closer proximity would reduce the required diffusion distance as well as increase the local concentration of enzymes and substrates, thereby increasing probabilistic processing by the second enzyme (Figure 5a, Figure 5b); this is often put forward as the main rationale for spatially organising enzymes using synthetic scaffolds and compartments^11,48–52^. However, it is possible that extreme proximity can result in unfavourable relative orientations of active sites that hinders reactions^53^. Translationally fused green and red fluorescent proteins were found to exhibit higher FRET when packaged inside bacteriophage P22 VLPs, indicating that the confinement forced the fused domains closer together^28^. For the Linked constructs (translationally fused FPPS-NES), we observe less linalool production upon encapsulation, which may be related to the constraints on enzyme orientation imposed by encapsulation and consequently less competition at the FPP node (Figure 5b). Compared to compartment-bounded enzyme configurations where enzyme spacing and positions are constrained by the VLP shell, Linked Free could also conceivably form enzyme clusters of a different density and distribution of FPPS and NES due to their multimerism (Figure 5c). It has been proposed that scaffolded or fused enzymes could form a network with adjacent proteins via enzyme homo-oligomerisation^11,51^, producing regions with high local concentrations of enzymes. In one study, formation of these large *in vivo* clusters was reported to be essential for a ‘metabolic channelling’ effect to occur, while enzyme fusion alone did not have a sizeable impact^54^.

**Figure 5.**
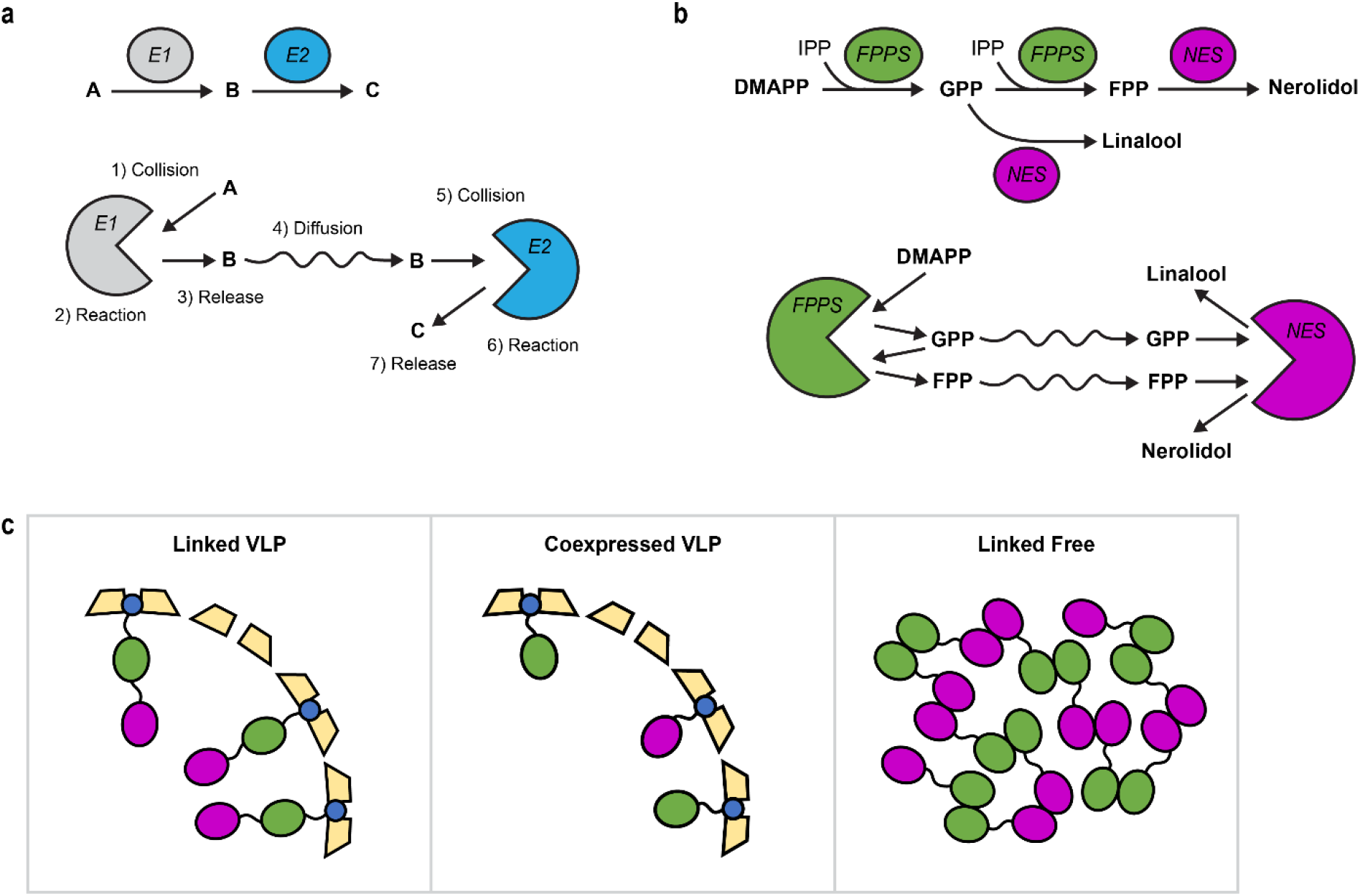
Proposed effects of enzyme spatial organisation on substrate accessibility. (a) In a simplified model of a two-step reaction where the conversion of substrate ‘A’ into final product ‘C’ proceeds via intermediate ‘B’, molecules of B have to diffuse from the enzyme that catalyses the first reaction (E1) towards the second enzyme in the cascade (E2) before the next step can proceed. (b) FPPS catalyses the two-step conversion of DMAPP into FPP, which proceeds *via* the intermediate GPP. NES is a promiscuous enzyme that can convert GPP into linalool or FPP into nerolidol. Any GPP that is released by FPPS could potentially be consumed by NES before it can be catalysed by FPPS into FPP. A small average distance between FPPS and NES therefore favours linalool production, while a larger distance favours nerolidol production. For simplicity, the co-substrate IPP is not shown in the cartoon. (c) The average inter-enzyme distance and substrate accessibility is expected to differ between the Linked VLP, Coexpressed VLP, and Linked Free configurations. In the case of Linked Free, dense enzyme clusters could potentially be formed by enzyme homo-oligomerisation.

However, increased inter-enzyme proximity alone is often insufficient for explaining the behaviour of spatially organised enzymes in biocatalytic cascades, especially given that most enzymes are not diffusion-limited^51,55^. More recently, there has been increased appreciation around additional factors that can affect enzyme kinetics such as active site orientation^53,56,57^, macromolecular crowding^52,58,59^, and physicochemical properties of the scaffold or compartment itself^60–63^. There are still many unknowns around how self-assembling protein compartments modulate the behaviour of native and non-native enzymes. The co-encapsulation of enzyme cascades in self-assembling protein compartmentalisation often produces idiosyncratic effects that are platform- and enzyme-dependent, reflecting our limited understanding of such systems. Elucidating the possible roles of VLP architecture on enzyme properties is not only relevant to recombinant protein compartments, but also contributes to the understanding how natural self-assembling protein compartments work.

## CONCLUSIONS

We explored two parallel approaches for *in vivo* protein co-encapsulation in yeast, demonstrating high cargo loading and effective co-localisation of fluorescent cargo proteins. Implementation of the compartment platform to co-encapsulate two successive enzymes in the nerolidol biosynthesis pathway greatly improved nerolidol titres compared to non-scaffolded enzymes, with the Linked VLP approach delivering a ∼30-fold increase and the Coexpressed VLP approach an ∼11-fold increase. This is also the first report of a self-assembling artificial yeast compartment enhancing the productivity of a metabolic pathway. Compartmentalisation was able to stabilise NES while being inherently more flexible than the enzyme fusion approach, potentially enabling the titration of enzyme local concentration and stoichiometry. In metabolic engineering studies employing a programmable synthetic scaffold, the stoichiometry of co-localised enzymes was repeatedly found to be an important parameter for optimising productivity^3,64–66^. In the MPyV VLP platform, tuning of the ratio of co-encapsulated enzymes is possible with the Coexpressed VLP strategy, using expression promoters of different strengths.

A surprising observation from this work was the influence of enzyme spatial organisation on the product profile, specifically the nerolidol:linalool ratio. This has industrial potential for obtaining high purity product directly from the fermentation, as illustrated by Coexpressed VLP (wtFPPS) that greatly enhanced nerolidol production but produces virtually no linalool. Based on this finding, it would be useful to explore if adjusting the ratio of encapsulated FPPS:NES could be exploited to drive the product preference towards a specific direction. More broadly, we speculate that spatial organisation using self-assembling compartments could be a new route for controlling the product specificity of promiscuous enzymes.

The modularity of the isoprenoid pathway means lessons from this study might be applicable to the bioproduction of diverse isoprenoids. It would be interesting to explore if the titre enhancement and product specificity observed here are reproducible with other *in vivo* compartment types. In addition, a detailed understanding of the mechanism of product specificity can only be elucidated by performing further *in vitro* and *in silico* enzyme studies. This work highlights the importance of exploring multiple scaffolding strategies when evaluating a biocatalytic cascade, and further establishes the modular MPyV platform as a useful tool for organising and stabilising metabolic enzymes *in vivo*.

## METHODS

### Molecular cloning and strain generation

All cloning was performed by isothermal assembly (NEBuilder HiFi DNA Assembly Master Mix, NEB #E2621), using the plasmid ΔVP1 + VP2C-GFP^14^ as the backbone vector. The GFP variant used for this study was yeast-enhanced GFP (yeGFP3^67^/GFPmut3^68^). The mRuby3-only VLP control (P_GAL1_-ΔVP1 + P_GAL10_-VP2C-mRuby3) was generated by directly replacing the GFP sequence with that of mRuby3^69^. The GFP gene was excised from the vector by double digesting with BamHI and BglII followed by gel purification of the backbone. The sequence for mRuby3 was PCR-amplified from a synthetic gene (synthesized by Integrated DNA Technologies Inc.) to add the appropriate 5’ and 3’ overlapping sequences and assembled together with the linearised backbone vector. For fusion constructs (containing GFP-mRuby3 or FPPS-NES), dsDNA fragments were generated of each fusion partner with a short connecting linker peptide (RSAGGGGTGGAEL) added using PCR primer overhangs. The two fragments were then co-assembled with the linearised backbone vector. For the Coexpressed construct (P_GAL1_-ΔVP1 + P_GAL10_-VP2C-GFP + P_GAL7_-VP2C-mRuby3), VP2C-GFP and T_GAL10_-P_GAL7_ fragments were amplified from the GFP VLP plasmid and yeast genomic DNA respectively. The two fragments were then co-assembled with the mRuby3-only VLP plasmid linearised by digestion with ClaI. The Linked VLP and Coexpressed VLP constructs for nerolidol production were cloned using similar strategies. The FPPS(F96W-N127W) constructs were generated by excising wtFPPS from each corresponding NES cassette by double digesting with BamHI and BglII, and then cloning in a FPPS(F96W-N127W) fragment with the appropriate overhangs. Yeast strain genotype details, PCR primers, and synthetic gene sequences are provided in Table S1, Table S2, and Table S3 respectively. The isothermal assembly and yeast transformation procedures are as previously reported^14^. For nerolidol bioproduction experiments, 3 individual transformed colonies from each construct were maintained as biological replicates. Strains were cultured overnight in YPD medium and stored as 20% v/v glycerol stocks at -80 °C.

### VLP expression and purification

GFP and mRuby3 particles were expressed and purified as previously described^14^, with an additional step prior to polishing by size-exclusion chromatography: mRuby3-containing VLP samples (collected after iodixanol cushion ultracentrifugation) were incubated at 37 °C for ∼15 hours to allow complete fluorophore maturation of mRuby3. Colour change of the samples from colourless/pale yellow to pink could be seen by eye.

### VLP characterisation

Transmission electron microscopy (TEM), nanoparticle tracking analysis (NTA), SDS-PAGE, and native agarose gel electrophoresis were performed essentially as described in our previous work^14^. For SDS-PAGE and native agarose gel electrophoresis, 3 μg purified VLP sample was loaded in each lane. For fluorescence imaging of the native agarose gel using a ChemiDoc MP Imaging System (Bio-Rad), the ‘Fluorescein’ preset was used for GFP (blue epi excitation, 530/30 nm filter) and the ‘Cy3’ preset for mRuby3 (green epi excitation, 605/50 nm filter).

### Förster resonance energy transfer (FRET) assay

Each sample consists of 3 technical replicates (200 μl each) in black 96-well microtitre plates. Purified VLPs were diluted in Buffer A (20 mM MOPS, 150 mM NaCl, 1 mM CaCl, pH 7.8) to ∼10 μg/ml and measured with a microplate reader (Tecan Infinite 200 Pro M Plex). Settings: Excitation wavelength = 450 nm (10 nm bandwidth), emission wavelength = 485-650 nm (step size = 5 nm), gain = 150, number of flashes = 10, integration time = 50 μs.

### Superresolution structured illumination microscopy (SR-SIM)

Glass-bottom dishes (MatTek #P35G-1.5-14-C) were covered with ∼250 μl 0.1% w/v poly-L-lysine solution (ProSciTech Pty Ltd #C500) and left to air dry overnight at room temperature. The coated surface was washed thoroughly with distilled water to remove excess poly-L-lysine and air-dried again. Purified VLP samples were diluted to ∼1 μg/ml in glycerol buffer (90% v/v glycerol, 20 mM Tris-Cl, pH 8.4). 200-300 μl diluted sample was carefully settled on the coverslip, tilting the dish slightly to allow the viscous sample to flow and create an even layer. Dishes were prepared at least a few hours to a few days before imaging (stored in the dark at 4 °C) to allow time for the particles to adhere to the coverslip.

Imaging was performed with a Zeiss ELYRA PS.1 equipped with an alpha plan-apochromat 100×/NA 1.46 oil immersion objective and an imaging chamber set at 30 °C. Acquisition settings: 3D SR-SIM mode (11 slices, 0.101 μm interval); 5 rotations per slice; 1280×1280 px; 488 nm excitation/495-550 nm emission, 30% laser, 100 ms exposure (GFP); 561 nm excitation/570-620 nm emission, 30% laser, 100 ms exposure (mRuby3). After placing each new dish, temperature was allowed to re-equilibriate for several minutes before imaging. Sample focusing was performed using the GFP channel, working as fast as possible and moving to a new spot every time to minimise photobleaching. A large z-interval was required to cover the full range of both colour channels in order to account for chromatic aberration and slight variations in the glass surface. SR-SIM image reconstruction settings: Noise filter = -6, Baseline = shifted, Use raw scale (ScaleToFit = False), PSF = Theoretical, Sectioning = 100, 83, 83. Channel alignment was then applied individually to every processed image (only in Linked and Coexpressed datasets – the mixed samples did not have enough signal overlap to enable automatic alignment). Alignment settings: Fit, Affine. The dataset contains 4 images for Linked, 6 images for Coexpressed, and 5 images for the mixed control. The number of included particle ROIs is indicated in each plot. Reconstructed and aligned image files were batch analysed with ImageJ (v1.52n) macro scripts. The detailed analysis workflow and ImageJ macro scripts are provided in Supporting Information.

### Nerolidol/linalool production

A two-phase galactose fermentation protocol was used, using dodecane as an inert organic overlay for trapping nerolidol, linalool, and other volatile isoprenoid pathway metabolites. Strains were pre-cultured overnight in YPD medium and inoculated at OD_600_ = 0.05 into 20 ml of a galactose-containing rich medium (2% w/v Bacto peptone, 1% w/v Bacto yeast extract, 2% w/v galactose, and 0.5% w/v glucose) to initiate the fermentation. The cultures were overlayed with 2 ml sterile dodecane. The methods for flask fermentation, metabolite analysis by high-performance liquid chromatography (HPLC), and whole-cell proteomics were the same as previously^39^.

### Data analysis and plotting

Charts were plotted using the Matplotlib package in Python 3. Analysis of NTA data was performed using parameters described previously^14^. For FRET emission spectra and SR-SIM histograms, data was smoothed using a Savitzky-Golay filter using the following parameters: window size = 9, polynomial order = 3 for FRET; window size = 7, polynomial order = 3, binwidth = 0.05 for SIM.

## ASSOCIATED CONTENT

Supporting Information file (PDF).

Plasmids used in this work are available from Addgene – refer to Supporting Information file for details

## ACKNOWLEDGEMENTS

The work was performed on the traditional lands of the Yugarabul, Yuggera, Jagera and Turrbal peoples. This work was supported in part by a Research Grant from the Human Frontier Science Program (Ref.-No: RGP0012/2018) and a CSIRO-UQ strategic fund. L.C.C was supported by a Commonwealth Research Training Program scholarship and a CSIRO-UQ Postdoctoral Fellowship. Additional funding for this project was provided through a CSIRO Synthetic Biology Future Science Platform PhD Top-up Scholarship. F.S. and B.P. acknowledge support from CSIRO in the form of a Synthetic Biology Future Science Platform Fellowship. B.P. acknowledges support from the Australian Research Council Centre of Excellence in Synthetic Biology (project number CE200100029), which is funded by the Australian Government. Metabolomics Australia is supported by BioPlatforms Australia through the Commonwealth Government’s National Collaborative Research Infrastructure Strategy (NCRIS). SR-SIM imaging was performed at the Queensland Brain Institute’s Advanced Microscopy Facility, using equipment funded through the ARC LIEF grant LE100100074. The authors are grateful to Rumelo Amor and Sandrine Roy for technical assistance with SR-SIM and Mitchell J. O’Sullivan (Queensland University of Technology) for assistance with writing ImageJ and Python scripts for data analysis. The authors thank Gert Talbo (Metabolomics Australia, Queensland Node) for technical assistance with whole-cell proteomics. The authors acknowledge the facilities and technical assistance of the Microscopy Australia Facility at the Centre for Microscopy and Microanalysis, The University of Queensland.

## AUTHOR CONTRIBUTIONS

L.C.C., F.S., and C.E.V designed experiments. L.C.C. conducted experiments. L.L. performed whole-cell proteomic analysis by LC-MS/MS. M.R.P. performed metabolite analysis by HPLC. B.P. developed the base strain for nerolidol production and assisted with experimental design and data interpretation. Z.L. assisted with proteomics and metabolomics experiments. F.S., C.E.V., and G.S. supervised the project. L.C.C., F.S., C.E.V., and G.S. wrote the initial manuscript draft. All authors read and approved the final manuscript.

