## Supplementary Information for "Synthetic *in vivo* compartmentalisation improves metabolic flux and modulates the product profile of promiscuous enzymes"

* Corresponding authors:

**Supplementary Methods**


SIM data analysis workflow 3

Squalene and ergosterol extraction and quantification 6

Quantitative PCR for detecting multicopy gene integration 6

Cargo protein stabilisation assay 7

References for Supporting Information 24


**Supplementary Figures and Tables**


Figure S1. Assessing FRET assay parameters using single-fluorophore VLP controls. 8

Figure S2. SR-SIM image analysis. 9

Figure S3. Analysis of exported SR-SIM ROI data (before filtering). 10

Figure S4. TEM images of VLPs isolated from NES-expressing strains. 11

Figure S5. Levels of key isoprenoid pathway metabolites at 72 h p.i. (HPLC). 12

Figure S6. Whole-cell proteomics of wtFPPS-overexpressing strains. 13

Figure S7 Whole-cell proteomics of FPPS(F96W-N127W)-overexpressing strains. 14

Figure S8. Verification of multicopy gene integration by quantitative PCR. 15

Figure S9. VLP assembly is required for *in vivo* stabilisation of cargo proteins. 16

Table S1. Genotype details of *S. cerevisiae* strains used in this work. 17

Table S2. PCR primer sequences for strain construction. 19

Table S3. Synthetic gene sequences. 22

**SIM data analysis workflow**

Reconstructed and aligned image files were batch analysed with custom ImageJ (v1.52n) macro scripts. Each image is analysed independently. The workflow is as follows:

1. Create a maximum intensity projection image for each channel (flattens Z-stack into a single image).
2. Retrieve the representative image ‘background value’ for each channel. This is taken to be the (mean + 3 standard deviations), given that particles only occupy a small proportion of the total pixels. In our hands, (mean + 3 standard deviations) gave better removal of grid artifacts compared to using the mean alone. Due to unavoidable variability in SR-SIM image reconstruction, we were unable to use a fixed value for background subtraction across the whole dataset.
3. Subtract the background value from each channel.
4. Create a maximum intensity projection image by merging both channels. This creates a single-channel image containing all particle locations.
5. Generate ROIs (regions of interest) by thresholding the merged image. Threshold and ROI size parameters need to be manually optimised until particles are correctly selected. We recommend erring on the side of false positives for this step, as spurious ROIs can be discarded in downstream analysis. Export ROIs to file.
6. Use the generated ROI list to analyse the two-channel maximum intensity projection image (from step 1). Analyse each channel separately using the same ROIs. Export particle measurements to file.

Downstream analysis (performed in Excel and Python):

7. Combine particle data of the same sample to create the full dataset.
8. We chose to represent each ROI by the highest pixel value, in order to capture the maximum particle signal. ROI integrated density is another suitable measure.
9. Reproduce the background subtraction step by subtracting the lower threshold value from each ROI. For this step, we chose to use the mean as the threshold as it produced a more ‘even’ distribution compared to (mean + 3 standard deviations). This is done separately for each channel.
10. If necessary, set negative values to zero.
11. Sum Red and Green pixel values. Calculate the % Red signal over the combined signal.
12. If a large number of spurious ‘particles’ (grid artifacts) are present, remove ROIs with low signal from the dataset. Scatter plots of Green + Red values against % Red (Figure S3) are helpful to identify artifacts and determine a suitable cut-off. At low signal intensities, it becomes difficult to distinguish particle fluorescence from SR-SIM reconstruction artifacts. In addition, the dimmest particles are the most susceptible to the effects of photobleaching and focal plane mismatch. Taking these factors into account, we chose to exclude dim ROIs from further analysis (defined here as Green + Red < 350 a.u.) (see Figure S3 for all particle data).

ImageJ (v1.52n) macro scripts:

**MaxprojectROI.ijm**

```
// This script is run first
// First enable Bio-Formats Windowless Importer before running script

function processFile(input, output, filename)
{
    // Open SIM image file in .czi format (from Zeiss ELYRA PS.1)
    open(input + File.separator + filename);

    // Create a maximum intensity projection image
    run("Z Project...", "projection=[Max Intensity]");
    saveAs("Tiff", output + File.separator + "MAX_" + filename);

    // Determine the signal cutoff values separately for each channel
    Stack.setChannel(1); // Green channel
    run("Measure");
    meanG = getResult("Mean");
    stdevG = getResult("StdDev");

    // Use Mean + 3 STD as the background signal cutoff value. This gives 'cleaner' removal of grid artifacts compared to using the mean.
    cutoffG = meanG + 3*stdevG;

    // Saturate a small percentage of the brightest pixels to remove outliers
    run("Enhance Contrast...", "saturated=0.001");

    // Retrieve the 99.9995th percentile pixel value to use as the upper cutoff
    getMinAndMax(min, maxG);

    print(filename + ", mean, stdev, cutoff, max99.9995");
    print("Green:, " + meanG + "," + stdevG + "," + cutoffG + "," + maxG);

    // Rescale image display based on cutoff values
    setMinAndMax(cutoffG, maxG);
    run("Apply LUT","slice");

    // Repeat for the next channel
    Stack.setChannel(2); // Red channel
    run("Measure");
    meanR = getResult("Mean");
    stdevR = getResult("StdDev");
    cutoffR = meanR + 3*stdevR;
    run("Enhance Contrast...", "saturated=0.001");
    getMinAndMax(min, maxR);
    print(filename + " Red: mean, stdev, cutoff, max99.9995");
    print("Red:, " + meanR + "," + stdevR + "," + cutoffR + "," + maxR);
    setMinAndMax(cutoffR, maxR);
    run("Apply LUT","slice");

    // Merge the maximum intensity projections from both channels to generate an ROI map
    run("Z Project...", "projection=[Max Intensity]");
    saveAs("Tiff", output + File.separator + "merge_" + filename);
    run("Clear Results");

    // Use the thresholding method to segment ROIs from background
    setThreshold(4000, 65535); // Values are determined on a case-by-case basis
    run("Analyze Particles...", "size=0.002-0.08 exclude add"); // Parameters determined on a case-by-case basis
    roiManager("Save", output + File.separator + filename + "RoiSet.zip");

    wait(2000); // Allows time to manually inspect ROIs (in ms)
```

```
    run("Close");
    close("*");
}

// Create an new output folder beforehand, then specify the input and output directories here
// EDIT THIS: Folder with .czi image files
inputDir = "C:/Users/Username/Desktop/Examplefolder1";
// EDIT THIS: Empty folder for output images and data
outputDir = "C:/Users/Username/Desktop/Examplefolder2";

list = getFileList(inputDir);
for (i = 0; i < list.length; i++)
    if(startsWith(list[i], "ExampleName")) // EDIT THIS: Optional step for testing script on a subset of images
        if(endsWith(list[i], ".czi"))
           processFile(inputDir, outputDir, list[i]);

selectWindow("Log");
saveAs("Text", outputDir + File.separator + "log" + ".csv")
```

### Exportparticles.ijm

```
// This script is run only after running MaxprojectROI.ijm

function processFile(input, output, filename)
{
    // Identify all maximum intensity projection files
    maxFilename = replace(filename, "MAX_", "");

    // Identify all ROI data files
    roiFilename = replace(maxFilename, ".tif", ".cziRoiSet.zip");
    open(input + File.separator + filename);

    Stack.setChannel(1);
    roiManager("Open", input + File.separator + roiFilename);
    roiManager("Measure");
    saveAs("Results", output + File.separator + filename + "MeasureG.csv");
    run("Clear Results");

    Stack.setChannel(2);
    roiManager("Measure");
    saveAs("Results", output + File.separator + filename + "MeasureR.csv");
    close("*");
    close("Results");
    close("ROI Manager");
}

// EDIT THIS: Specify input and output directories
inputDir = "C:/Users/Username/Desktop/Examplefolder1";
outputDir = "C:/Users/Username/Desktop/Examplefolder2";

list = getFileList(inputDir);
for (i = 0; i < list.length; i++)
    if(startsWith(list[i], "ExampleName")) // EDIT THIS: Optional step for testing script on a subset of images
        if(endsWith(list[i], ".tif"))
            processFile(inputDir, outputDir, list[i])
```

**Squalene and ergosterol extraction and quantification**

Squalene and ergosterol were extracted and quantified by HPLC as previously described[1]. For each strain, one tube of stored cell samples was used for the extraction while a separate tube was used to determine the dry cell mass. Frozen cell pellets (from 2 ml culture at 72h p.i.) were thawed and resuspended in 400 µl 2M KOH in methanol. 100 µl internal standard solution (400 µM pyrene, 10 g/L pyrogallol in methanol) was also added to each sample. Samples were incubated at 80 °C in 2.2 ml screw-capped tubes for 1 hour, with thorough vortexing every 15 min. 500 µl distilled water and 1 ml hexane was added to the mixture and lipids were extracted by vigorous vortexing for 30 min. The upper 750 µl of the hexane phase was evaporated with a vacuum concentrator, and the pellet resuspended in 150 µl 1:1 methanol:ethanol with 2 g/L pyrogallol. 20 µl from each sample was injected through a Gemini C18 column (150×4.6 mm, 3 µm, 110 Å, PN: 00F-4439-E0) with a guard column (SecurityGuard Gemini C18, PN: AJO-7597). Analytes were eluted isocratically at 1 ml/min with methanol at 35 °C. Analytes were monitored using a diode array detector at the following UV wavelengths: 200 nm (for squalene), 274 nm (for ergosterol), and 334 nm (for pyrene).

**Quantitative PCR for detecting multicopy gene integration**

Quantitative PCR (qPCR) assays were performed on yeast genomic DNA to measure the copy number of the 4 genes in the nerolidol expression cassette (ΔVP1, VP2C-FPPS, NES, *K. lactis* URA3) relative to a single-copy housekeeping gene ACT1. All yeast strains in this work are haploid.

Yeast glycerol stocks were recovered on uracil drop-out plates and cultured in YPD liquid medium. Genomic DNA was extracted using the YeaStar Genomic DNA Kit (Zymo Research, #D2002) according to the manufacturer’s protocol. 20 µl qPCR reactions were set up using a SensiFAST SYBR Hi-ROX Kit (Bioline, #BIO-92005), containing 20 ng of genomic DNA and 400 nM of each qPCR primer. Thin-wall white polypropylene PCR tubes (Bio-Rad #TLS0851XTU) and ultraclear optical flat caps (Bio-Rad #35TCS0803) were used. VP2C-FPPS primers were designed to span VP2C, the flexible fusion linker, and *S. cerevisiae* FPPS (ERG20). The KlURA3 primers were designed to be specific to *K. lactis* URA3; these primers were found not to cross-detect *S. cerevisiae* URA3 (the base strain contains a non-functional ura3-52 allele).

Quantitative PCR primer sequences were as follows:

| **Primer name** | **Target gene** | **Sequence (5' - 3')** |
|---|---|---|
| VP1_qpcr_fw | MPyV ΔVP1 | GCACAAAGGCTTGCCCTAGACC |
| VP1_qpcr_rv | MPyV ΔVP1 | CTCTTGACCAACCGTAATATTGACCACC |
| VP2C_qpcr_fw | VP2C-FPPS | TTTAGGGAAAACGTTCAAGAATCTCTCTCTCC |
| ERG20_qpcr_rv | VP2C-FPPS | AAGAGTTACTCCAGATTGGATGTTGCC |
| AcNES1_qpcr_fw | NES | CTTGGGTTCAGCCAAAGATGAAGACC |

| | | |
|---|---|---|
| AcNES1_qpcr_rv | NES | CCTTATTCAAACATTTCCATGCTTCGG |
| klURA3_qpcr_fw | *K. lactis* URA3 | CAATTTGTGTGCTTCTCTTGACGTTCG |
| klURA3_qpcr_rv | *K. lactis* URA3 | TGCTTTCAATGGAACGACAGTACCC |
| ACT1_qpcr_fw | *S. cerevisiae* ACT1 | ACCAAGGTATCATGGTCGGTATGG |
| ACT1_qpcr_rv | *S. cerevisiae* ACT1 | GGTGTTCTTCTGGGGCAACTCTC |

Data was acquired with a Bio-Rad CFX96 Real-Time PCR Detection System coupled to a Bio-Rad C1000 thermal cycler, with the following protocol:

95.0°C for 3:00
95.0°C for 0:05
60.0°C for 0:30
Plate Read
GOTO 2, 39 more times
Melt Curve 65.0°C to 95.0°C : Increment 0.5°C for 0:05
Plate Read
10.0°C for 5:00

The experiment was performed with technical duplicates. Data was analysed using Bio-Rad CFX Manager software, with the cycle threshold (Ct) manually set at 1500 a.u. for SYBR Green. Relative gene expression levels were calculated using the standard delta-delta Ct method ($2^{-\Delta\Delta Ct}$)[2], with ACT1 as the internal reference gene. Bar chart values are the means of the technical duplicates.

**Cargo protein stabilisation assay**

We have previously developed a method for observing cargo protein stabilisation *in vivo* by MPyV VLPs, using destabilised GFP ($GFP_{Deg}$) as the model cargo[3]. In the current work, we used the same approach to determine if VLP assembly is essential for the stabilisation of VP2C-tagged cargo proteins (Figure S9). The method for flow cytometry and ultracentrifuge analysis is as previously reported, except the cultures were inoculated at starting $OD_{600}$ = 0.2 and were grown for 30h.

For the anti-VP1 dot blot, 2 μl of each sample collected after the ultracentrifugation step was spotted directly on a dry nitrocellulose membrane pre-marked with pencil. After air-drying for several minutes, the membrane was immersed in western blot transfer buffer (25 mM Tris, 192 mM glycine, 0.1% SDS, 20% v/v methanol) for 2 min. The blot was immediately staind with 0.1% w/v Ponceau S in 5% v/v acetic acid to check protein transfer. The subsequent blocking, antibody incubation, and imaging steps are as previously reported[3].

**Figure S1. Assessing FRET assay parameters using single-fluorophore VLP controls.**

**(a)** No significant emission in the FRET acceptor emission range was observed with GFP-only or mRuby3-only controls at 450 nm excitation. Values were blank-subtracted.

**(b)** A mixture of GFP-only and mRuby3-only VLPs produces an emission spectrum that is equivalent to the sum of their signals measured separately. This shows that GFP and mRuby3 in separate VLPs are unable to interact by FRET. The settings used here were the same as in the main text (450 nm excitation). A high concentration of mRuby3-only VLPs was used to obtain a clear emission peak. Values were blank-subtracted and is the mean of 3 technical replicates.

**a**

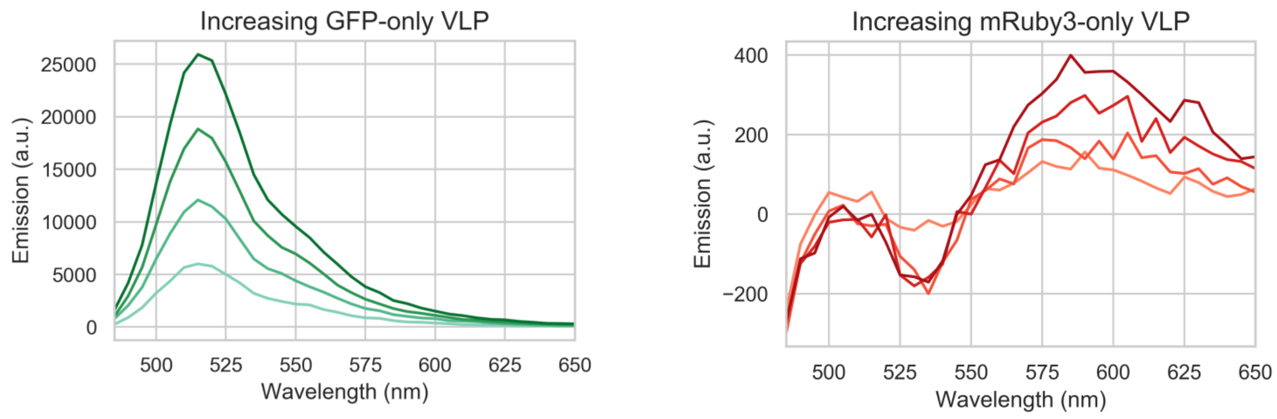


**b**

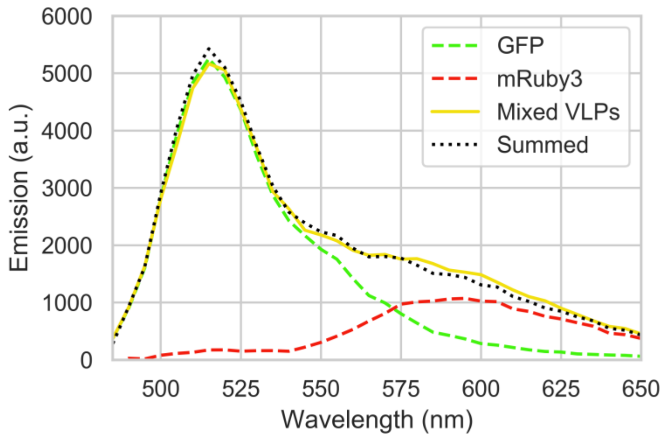

**Figure S2. SR-SIM image analysis.**

**(a)** A full uncropped two-channel SR-SIM image (of Linked VLPs) after image reconstruction, maximum intensity projection, and channel alignment (2430×2430 pixels, 49.36×49.36 µm). As in the main text, the GFP channel is displayed as green and mRuby3 channel displayed as magenta. In this sample image, the lower threshold value obtained from the ImageJ script (= mean pixel value) was set to black to match the final outcome of the analysis workflow.

**(b)** The pixel values of the two colour channels from (a) were summed to create a single-channel merged image (displayed in black and white here). This new output image would be further processed and analysed to generate a list of ROIs corresponding to individual ‘spots’.

**(c)** Zoomed view of a small region from (a).

**(d)** The same region from (c) overlaid with ROIs generated using the analysis workflow. The ROIs would be analysed using the ‘Measure’ tool and the resulting values exported. Further data filtering (post-export) was used to remove spurious ROIs and low-quality data (see **Figure S3**).

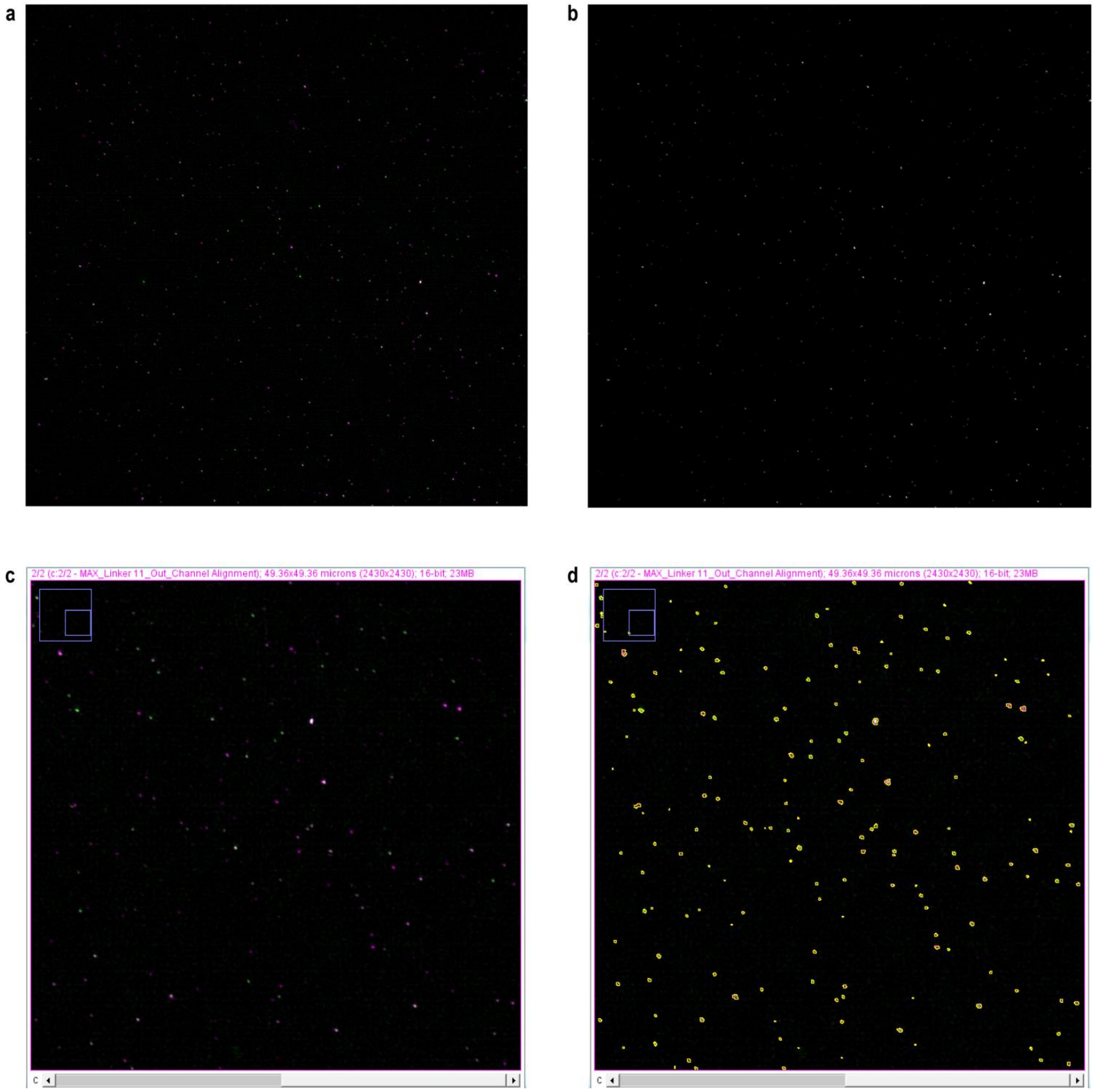

**Figure S3. Analysis of exported SR-SIM ROI data (before filtering).**

The dataset of all ROIs (= individual particles), including spurious ‘particles’ from SIM reconstruction artifacts. The number of particles in each plot (n) is shown as an inset.

**(a)** Scatter plots of the Green and Red signal values of each individual ROI.

**(b)** Sum of Green + Red signals plotted against % Red signal (= mRuby3 contribution). The dashed grey line shows the chosen lower cut-off for excluding ‘dim’ particles (Green + Red < 350 a.u.). The over-representation of high % Red particles in the region below the cut-off threshold is most likely due to photobleaching of GFP during imaging, which is much less photostable than mRuby3.

**(c)** Co-localisation analysis histograms.

For each set of scatter plots in (a) and (b), the same axis scales are used across samples to facilitate comparisons.

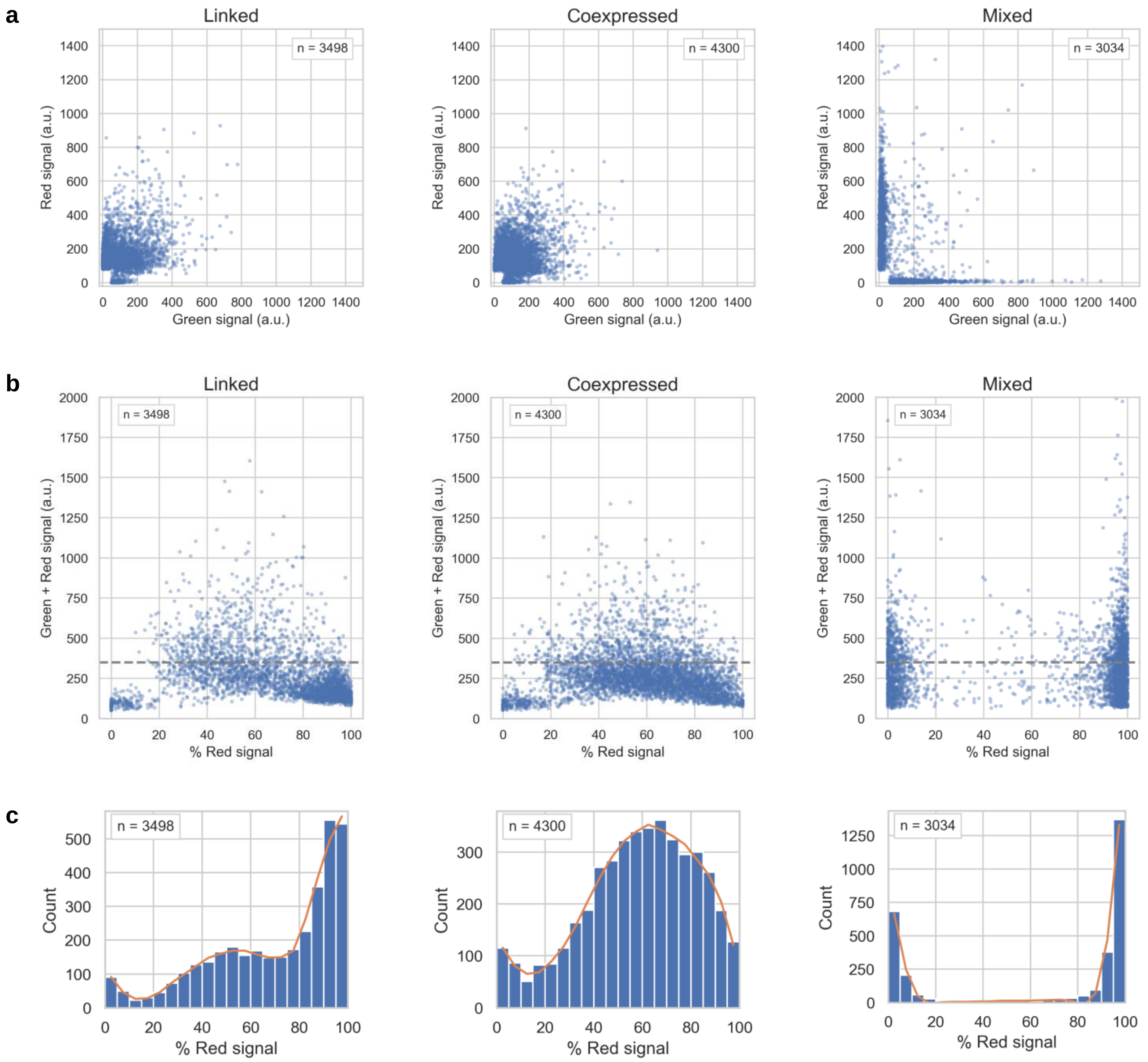

**Figure S4. TEM images of VLPs isolated from NES-expressing strains.**

Samples were isolated from 250 ml of yeast culture grown in a galactose-containing rich medium (1% w/v yeast extract, 2% w/v peptone, 2% w/v galactose, 0.5% w/v glucose). Cultures were inoculated at $OD_{600}$~0.2 and grown with a 20 ml dodecane overlay for 48 hours at 30 °C, 200 rpm shaking. Cell lysis, PEG8000-NaCl precipitation, ultracentrifugation, and TEM negative staining were performed as previously described[3]. Poor VLP yields were obtained because dodecane appeared to interfere with the lysis and purification method.

**(a)** VLPs isolated from Linked VLP (wtFPPS-overexpressing).

**(b)** VLPs isolated from Coexpressed VLP (wtFPPS-overexpressing).

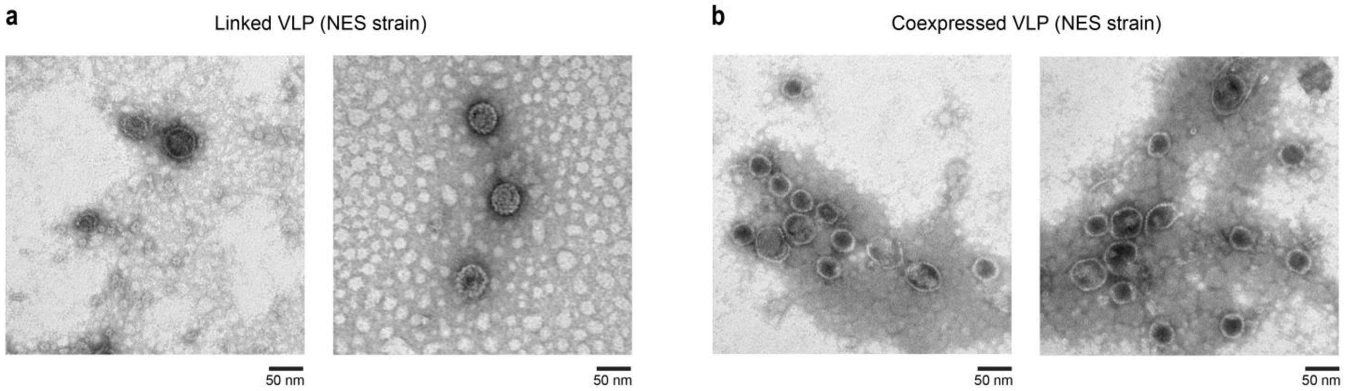

**Figure S5. Levels of key isoprenoid pathway metabolites at 72 h p.i. (HPLC).**

**(a)** Nerolidol and **(b)** linalool titres of strains overexpressing wtFPPS and NES (described in Figure 3 in the main text), presented as mg product per g dry cell mass (mg/g DCM). The nerolidol titre fold change relative to Coexpressed Free is indicated in red text.

**(c)** Concentrations of key ergosterol pathway metabolites squalene and ergosterol of strains overexpressing wtFPPS and NES, presented as mg per g dry cell mass (mg/g DCM).

Concentrations of isoprenoid pathway side products **(d)** farnesol and **(e)** geranylgeraniol of strains overexpressing wtFPPS and NES, presented as mg product per L cell culture (mg/L).

**(f)** Concentration of farnesol of strains overexpressing FPPS(F96W-N127W) and NES, presented as mg product per L cell culture (mg/L).

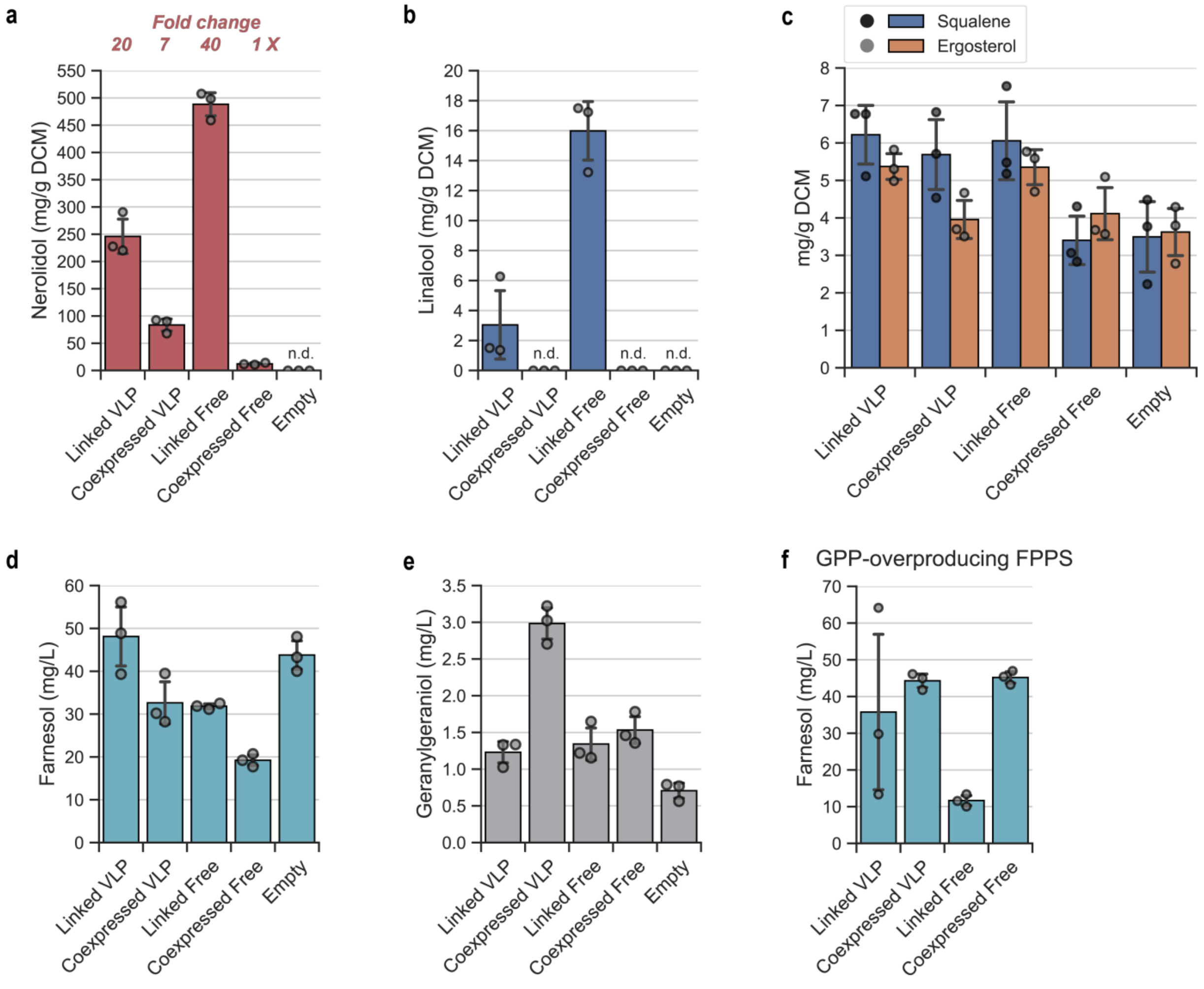

**Figure S6. Whole-cell proteomics of wtFPPS-overexpressing strains.**

'Normalised protein abundance' is a measure of the relative detected levels of each protein in the sample, normalised to the sum of all abundances for that sample. Since every protein behaves differently towards trypsinisation and ionisation, abundance comparisons can only be made for the same protein across different samples. In 'Scaled protein abundance', the level of each protein has been individually re-scaled so that the mean equals 100 across samples; this enables easier comparison especially for proteins detected at low abundance. Arbitrary units (a.u.) are used for all charts. The mean +/- 1 standard deviation and individual data points (3 biological replicates each) are shown. All proteins shown were detected with high confidence.

**(a)** Engineered nerolidol pathway genes. 1 out of the 3 Linked VLP strains expressed unusually high levels of the 5 genes in the nerolidol cassette (ΔVP1, VP2C, FPPS, NES, *Kluyveromyces lactis* URA3); this strain was later found to contain two genomically integrated copies of the nerolidol expression cassette (Figure S8). Note that in this strain, the measured VP2C level is similar to that of Coexpressed VLP, which has 2 copies of VP2C. EfmvaE (*Enterococcus faecalis* acetoacetyl-CoA thiolase/HMG-CoA reductase), EfmvaS (*E. faecalis* HMG-CoA synthase), and SKP1-OsTIR1 (S-phase kinase-associated protein 1 fused to *Oryza sativa* auxin receptor) are heterologous genes present in the base strain o57BR[4]. EfmvaE and EfmvaS were expressed using GAL2 and GAL1 promoters respectively, and could be useful as reporters for the degree of galactose induction in each strain.

**(b)** Housekeeping proteins (ACT1, TEF1/TEF2, PGK1) and **(c)** native galactose-inducible proteins (GAL1, GAL2, GAL7, GAL10) are useful to provide information on the stage of growth and extent of galactose induction in each strain. In the engineered nerolidol pathway, GAL1, GAL10, and GAL7 promoters were used to drive expression of ΔVP1, VP2C-FPPS-NES / VP2C-FPPS / FPPS, and VP2C-NES / NES respectively.

**(d)** Levels of un-engineered ergosterol pathway genes.

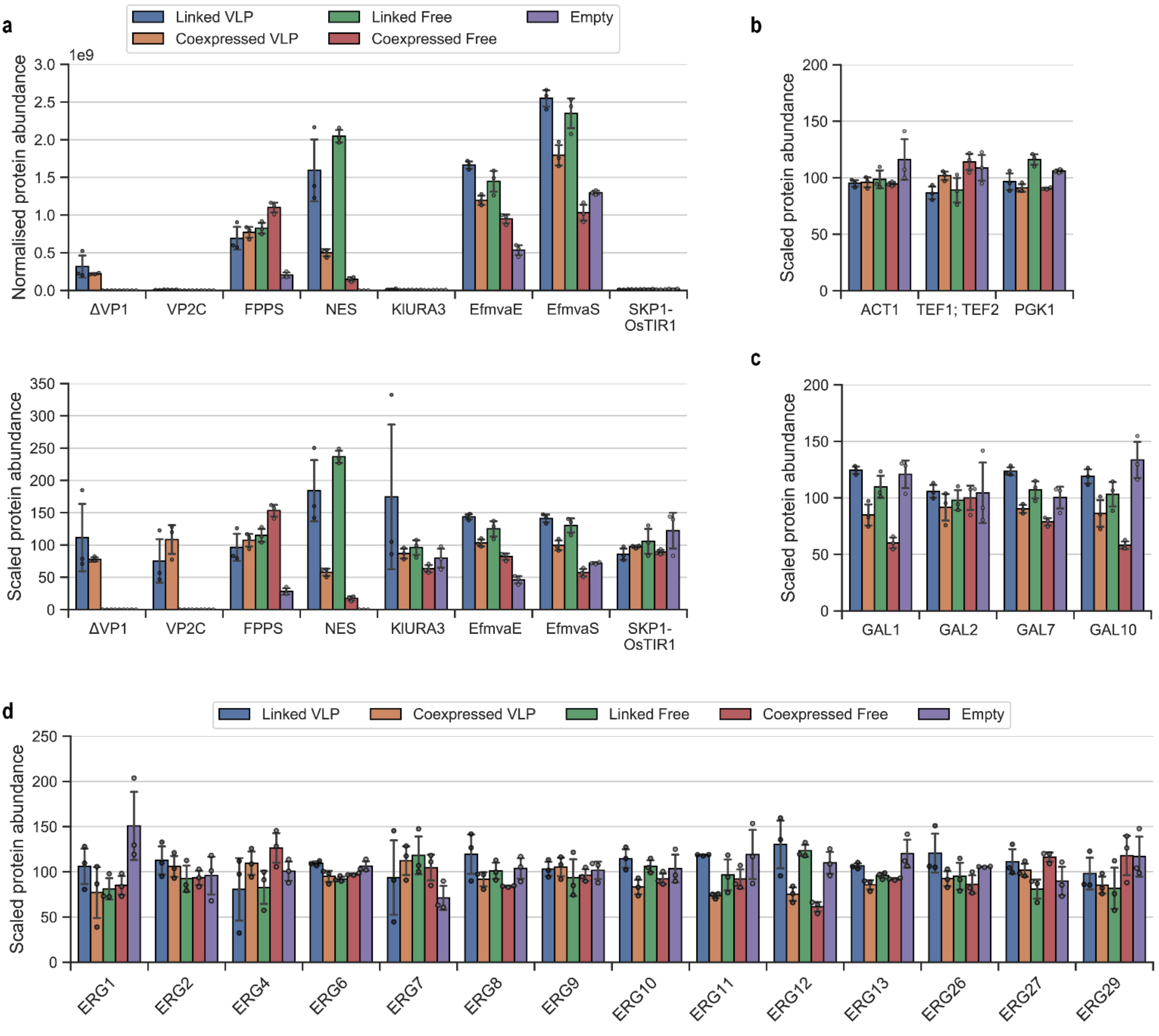

**Figure S7 Whole-cell proteomics of FPPS(F96W-N127W)-overexpressing strains.**

The level of each protein has been individually re-scaled so that the mean equals 100 across samples. Arbitrary units (a.u.) are used for all charts. The mean +/- 1 standard deviation and individual data points (3 biological replicates each) are shown. All proteins shown were detected with high confidence.

**(a)** Engineered nerolidol pathway genes, including genes introduced when constructing the base strain. 'FPPS' includes both wtFPPS and FPPS(F96W-N127W), which are indistinguishable due to their high sequence identity.

**(b)** Housekeeping proteins (ACT1, TEF1, PGK1).

**(c)** Native galactose-inducible proteins (GAL1, GAL2, GAL7, GAL10). In the engineered nerolidol pathway, GAL1, GAL10, and GAL7 promoters were used to drive expression of ΔVP1, VP2C-FPPS(F96W-N127W)-NES / VP2C-FPPS(F96W-N127W) / FPPS(F96W-N127W), and VP2C-NES / NES respectively.

**(d)** Levels of un-engineered ergosterol pathway genes.

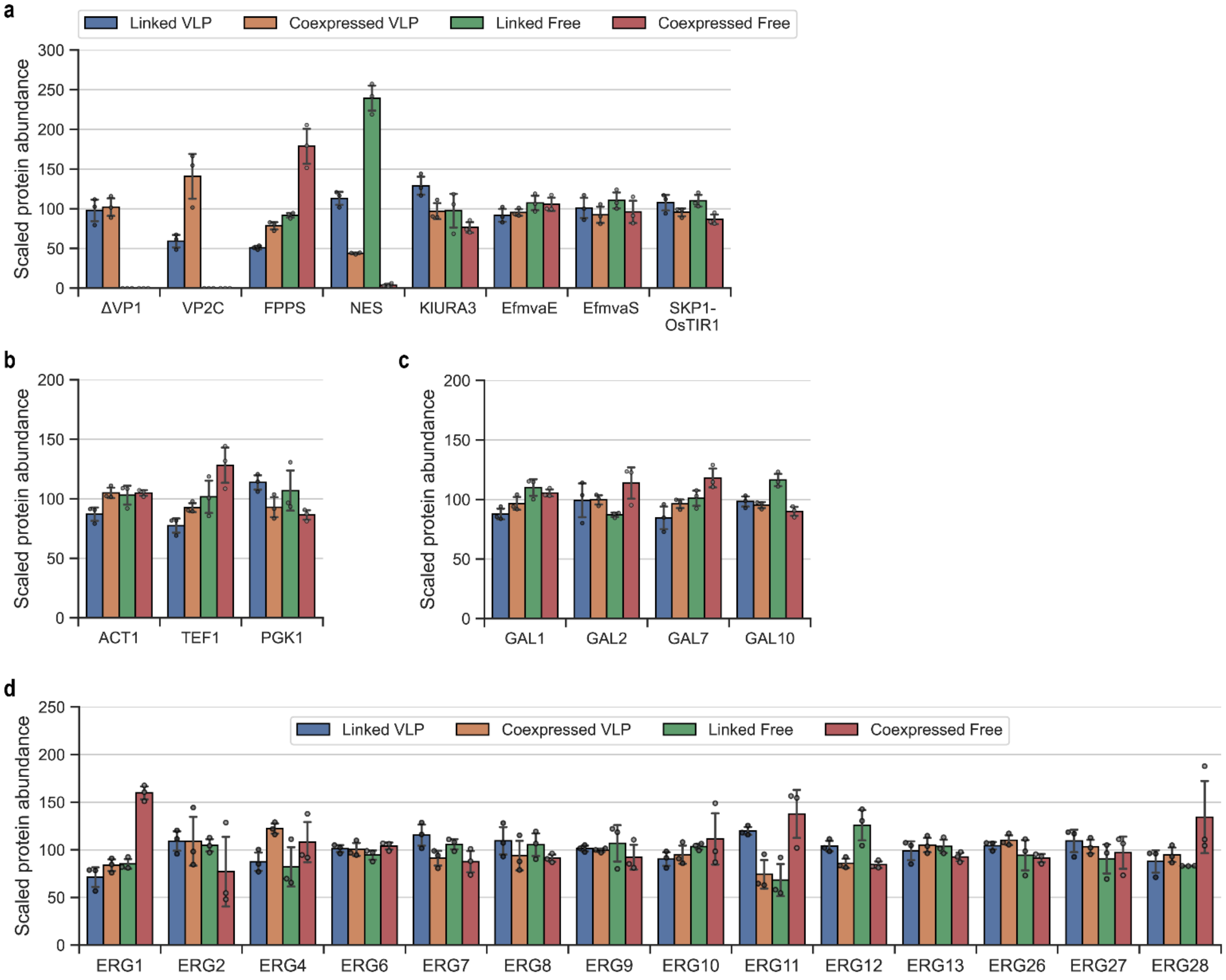

**Figure S8. Verification of multicopy gene integration by quantitative PCR.**

**(a)** Melt profiles and **(b)** agarose gel electrophoresis of PCR products were used to confirm the specificity and reproducibility of the assay.

**(c)** Gene copy number of wtFPPS-overexpressing strains (described in Figure 3), normalised to Linked VLP strain 1. The same biological replicate with unusually high protein expression in Figure 3 and Figure S6 (Linked VLP strain 2) has significantly increased copy numbers of all 4 nerolidol cassette genes.

**(d)** Due to high variability in nerolidol and linalool titres in Figure 4, the same analysis in (c) was performed on Linked VLP strains that overexpress FPPS(F96W-N127W). However, multicopy gene integration was not detected in this set. Gene copy numbers are normalised to Linked VLP strain 1.

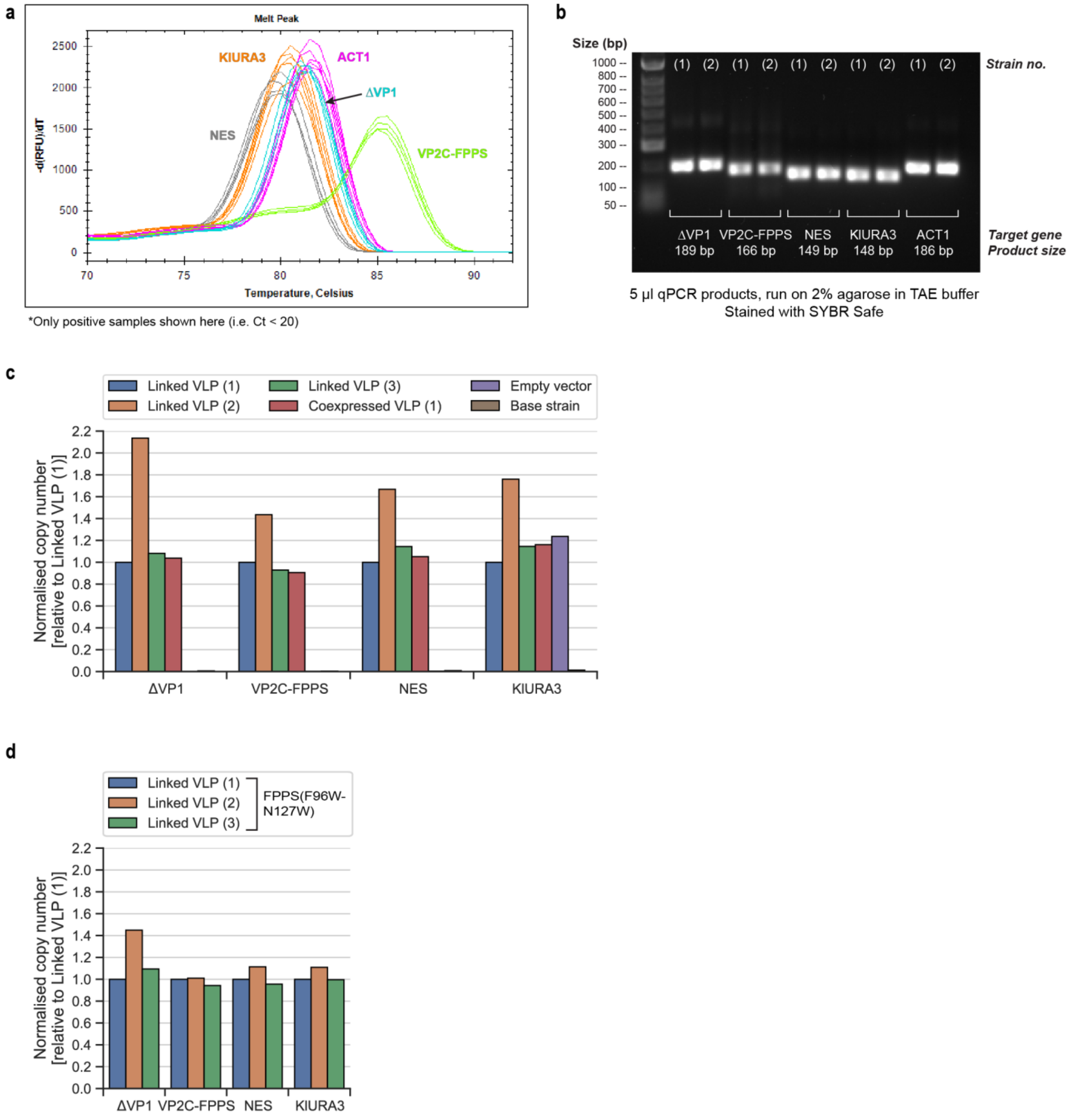

**Figure S9. VLP assembly is required for *in vivo* stabilisation of cargo proteins.**

MPyV VP1 missing its 63 C-terminal amino acids can form pentamers but cannot self-assemble into VLPs[5,6]. We performed this deletion on ΔVP1 to generate a ΔVP1(Δ63C) mutant, with controls expressing full-length ΔVP1 or without VP1.

**(a)** Flow cytometry for comparing GFP$_{Deg}$ fluorescence at 4 h and 30 h p.i.; distributions from 2 biological replicates are shown. Although binding with ΔVP1(Δ63C) appears to slightly dampen the destabilisation effect of the VP2C anchor, cargo sequestration by VLP assembly is essential for long-term stabilisation of GFP$_{Deg}$.

**(b)** Assembled VLPs were separated from other cellular proteins by ultracentrifugation through an iodixanol cushion. VLP-encapsulated GFP$_{Deg}$ presents as a fluorescent green band in the 'dense' layer, as can be seen in the ΔVP1 + VP2C-GFP$_{Deg}$ sample. Note that yeast cells are naturally autofluorescent in the GFP range, which explains the high apparent signal in the 'Free GFP$_{Deg}$' layer.

**(c)** 200 µl sample was collected from each of the 'Free GFP$_{Deg}$' and 'Encapsulated GFP$_{Deg}$' layers in (b) and imaged using the 'fluorescein' filter settings of a gel imager.

**(d)** Anti-VP1 dot blot to verify the absence of VLP assembly in the ΔVP1(Δ63C) mutant. 'Total protein' is a colorimetric image of the blot stained with Ponceau S. Both the 'Free GFP$_{Deg}$' and 'Encapsulated GFP$_{Deg}$' panels were imaged simultaneously on the same blot.

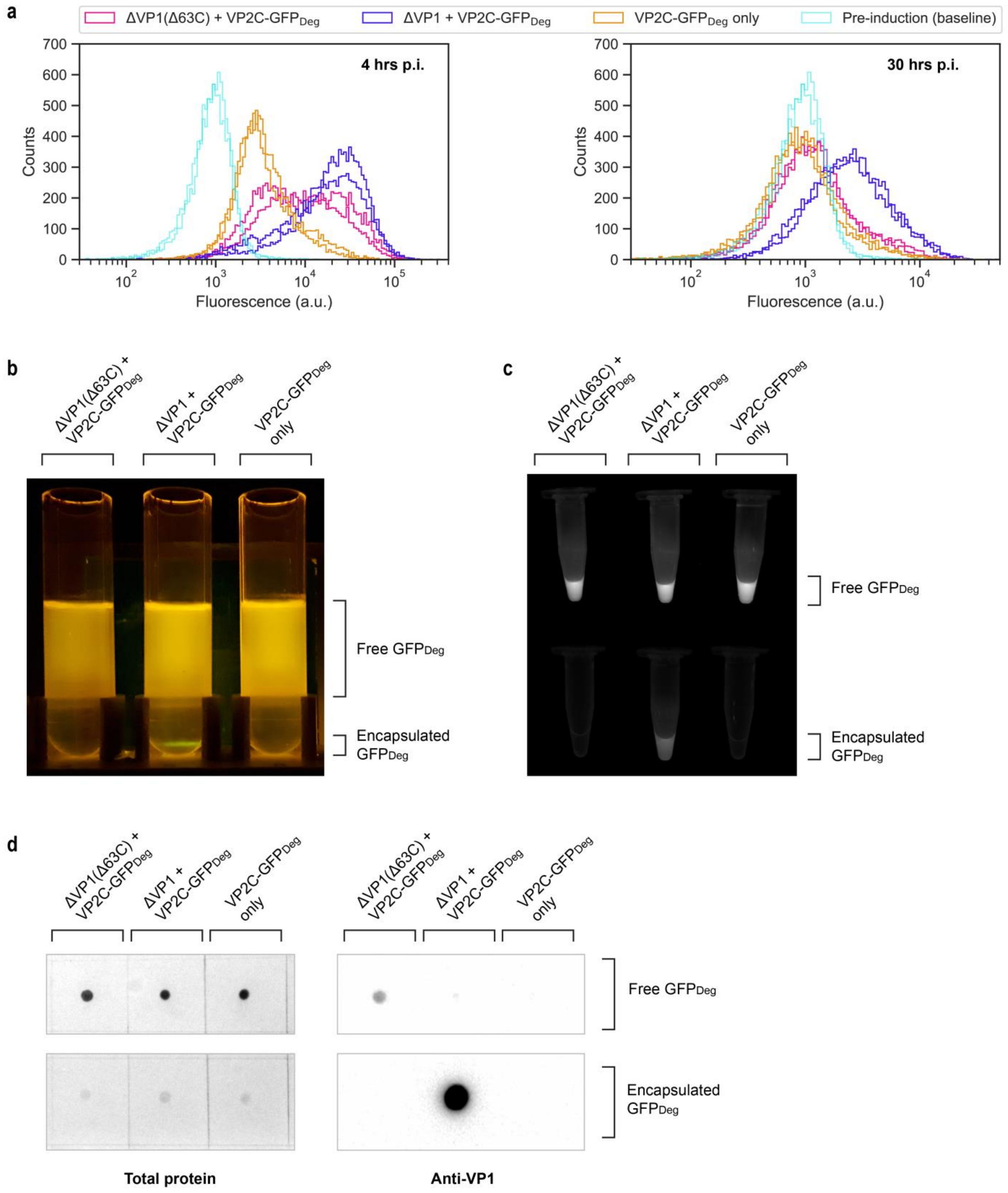

**Table S1. Genotype details of *S. cerevisiae* strains used in this work.**

Where available, the Addgene accession code for the yeast integrative plasmid (YIp) used to construct the strain is provided.

KlURA3 = URA3 gene (orotidine 5-phosphate decarboxylase) from *Kluyveromyces lactis.*

FPPS = ERG20 (yeast farnesyl diphosphate synthase), wild-type variant unless indicated otherwise.

| **Strain** | **Genotype** | **Description** | **Addgene #** | **Reference** |
|---|---|---|---|---|
| CEN.PK2-1C | MATa ura3-52 leu2-3,112 trp1-289 his3Δ1 MAL2-8c SUC2 | Base strain for VLP expression and mevalonate pathway overexpression strain o57BR | - | Entian and Kötter 2007[7] |
| GFP only | CEN.PK2-1C derivative;<br>ura3(1,704)::KlURA3-$T_{HIS5}$-(VP2C-GFP)-$P_{GAL10}$-$P_{GAL1}$-ΔVP1- $T_{URA3}$ | ΔVP1 + VP2C-GFP; for expressing VLPs for characterisation | 166676 | Cheah et al. 2021[3] |
| mRuby3 only | CEN.PK2-1C derivative;<br>ura3(1,704)::KlURA3-$T_{HIS5}$-(VP2C-mRuby3)-$P_{GAL10}$-$P_{GAL1}$-ΔVP1- $T_{URA3}$ | ΔVP1 + VP2C-mRuby3; for expressing VLPs for characterisation | 182429 | This work |
| Linked (GFP-mRuby3) | CEN.PK2-1C derivative;<br>ura3(1,704)::KlURA3-$T_{HIS5}$-(VP2C-GFP-mRuby3)-$P_{GAL10}$-$P_{GAL1}$-ΔVP1- $T_{URA3}$ | ΔVP1 + VP2C-GFP-mRuby3; for expressing VLPs for characterisation | 182430 | This work |
| Coexpressed (GFP + mRuby3) | CEN.PK2-1C derivative;<br>ura3(1,704)::KlURA3-$T_{HIS5}$-(VP2C-mRuby3)-$P_{GAL7}$-$T_{GAL10}$-(VP2C-GFP)- $P_{GAL10}$-$P_{GAL1}$-ΔVP1- $T_{URA3}$ | ΔVP1 + VP2C-GFP + VP2C-mRuby3; for expressing VLPs for characterisation | 182431 | This work |
| o57BR | CEN.PK2-1C derivative;<br>HMG2$^{K6R}$(-152,-1)::HIS3-$T_{EFM1}$-EfmvaS-$P_{GAL1}$-$P_{GAL10}$-ACS2-$T_{ACS2}$-$P_{GAL2}$-EfmvaE -$T_{EBS1}$-$P_{GAL7}$;<br>pdc5(-31,94)::$P_{GAL2}$-ERG12-$T_{NAT5}$-$P_{TEF2}$-ERG8-$T_{IDP1}$-$T_{PRM9}$-MVD1-$P_{ADH2}$-$T_{RPL15A}$-IDI1-$P_{TEF1}$-TRP1; ERG9(1333, 1335)::$T_{URA3}$-$P_{GAL7}$-MVD1-$T_{PRM9}$-$P_{GAL2}$-ERG12-$T_{NAT5}$-$T_{IDP1}$-ERG8-$P_{GAL10}$-$P_{GAL1}$-IDI1-$T_{RPL15A}$-$P_{ACS2}$-SKP1-OsTIR1;<br>MIG1(-1,1)::CUP1-AID* | Base strain for nerolidol production;<br>All mevalonate pathway enzymes overexpressed (except FPPS/ERG20) | - | Hayat et al. 2021[4] |
| Linked VLP | o57BR derivative;<br>ura3(1,704)::KlURA3-$T_{HIS5}$-(VP2C-FPPS-NES)-$P_{GAL10}$-$P_{GAL1}$-ΔVP1- $T_{URA3}$ | ΔVP1 + VP2C-FPPS-NES; nerolidol production strain | 182432 | This work |
| Coexpressed VLP | o57BR derivative;<br>ura3(1,704)::KlURA3-$T_{HIS5}$-(VP2C-NES)-$P_{GAL7}$-$T_{GAL10}$-(VP2C-FPPS)- $P_{GAL10}$-$P_{GAL1}$-ΔVP1- $T_{URA3}$ | ΔVP1 + VP2C-FPPS + VP2C-NES; nerolidol production strain | 182433 | This work |
| Linked Free | o57BR derivative; | VP2C-FPPS-NES; nerolidol | 182485 | Cheah et al. 2022[8] |

| | | | | |
|---|---|---|---|---|
| | ura3(1,704)::KlURA3-$T_{HIS5}$-(FPPS-NES)-$P_{GAL10}$-$P_{GAL1}$ | production strain | | |
| Coexpressed Free | o57BR derivative; ura3(1,704)::KlURA3-$T_{HIS5}$-NES-$P_{GAL7}$-$T_{GAL10}$-FPPS- $P_{GAL10}$ | FPPS + NES; nerolidol production strain | 182482 | Cheah et al. 2022[8] |
| Empty | o57BR derivative; ura3(1,704)::KlURA3 | Transformed with empty URA3 YIp; negative control | - | This work; plasmid pILGH4 from Peng et al. 2015[9] |
| Linked VLP (GPP-overproducing FPPS) | o57BR derivative; ura3(1,704)::KlURA3-$T_{HIS5}$-(VP2C-FPPS-NES)-$P_{GAL10}$-$P_{GAL1}$-$\Delta$VP1- $T_{URA3}$ | $\Delta$VP1 + VP2C-FPPS(F96W-N127W)-NES; nerolidol production strain | 182434 | This work |
| Coexpressed VLP (GPP-overproducing FPPS) | o57BR derivative; ura3(1,704)::KlURA3-$T_{HIS5}$-(VP2C-NES)-$P_{GAL7}$-$T_{GAL10}$-(VP2C-FPPS)- $P_{GAL10}$-$P_{GAL1}$-$\Delta$VP1- $T_{URA3}$ | $\Delta$VP1 + VP2C-FPPS(F96W-N127W) + VP2C-NES; nerolidol production strain | 182435 | This work |
| Linked Free (GPP-overproducing FPPS) | o57BR derivative; ura3(1,704)::KlURA3-$T_{HIS5}$-(FPPS-NES)-$P_{GAL10}$-$P_{GAL1}$ | VP2C-FPPS(F96W-N127W)-NES; nerolidol production strain | 182436 | This work |
| Coexpressed Free (GPP-overproducing FPPS) | o57BR derivative; ura3(1,704)::KlURA3-$T_{HIS5}$-NES-$P_{GAL7}$-$T_{GAL10}$-FPPS- $P_{GAL10}$ | FPPS(F96W-N127W) + NES; nerolidol production strain | 182437 | This work |

**Table S2. PCR primer sequences for strain construction.**

All gene constructs were cloned by isothermal assembly, which requires generating dsDNA fragments that overlap with adjacent fragments. Based on the desired order of promoters, genes, and terminators, fragments with the appropriate 5' and 3' overlaps can be produced by mixing and matching PCR primers with suitable overhangs.

| Name | Description | Sequence (5' - 3') | Length (nt) | PCR template | PCR fragment generated | Constructs |
|---|---|---|---|---|---|---|
| mRuby3_VP2C_fw | mRuby3 forward, with overhang for VP2C linker | GGCGGCGGTTCTATGGTGTCTAAGGGCGAAG | 31 | Synthetic mRuby3 gene | mRuby3 for VP2C | mRuby3-only VLP; Linked (ΔVP1 + VP2C-GFP-mRuby3) |
| mRuby3_AG4TGGA_fw | mRuby3 forward for GFP-mRuby3 fusion protein; appends a linker peptide (RSAGGGGTGGAEL) sequence | GCTGGTGGAGGAGGTACTGGTGGTGCTATGGTGTCTAAGGGCGAAG | 46 | Synthetic mRuby3 gene | mRuby3 for Linked | Linked (ΔVP1 + VP2C-GFP-mRuby3) |
| mRuby3_HIS5t_rv | mRuby3 reverse, with overhang for HIS5 terminator | ACTGTACATATACTGTTTAAATTAATCTATTTACTTGTACAGCTCGTCCATGC | 53 | Synthetic mRuby3 gene | mRuby3 | mRuby3-only VLP; Linked (ΔVP1 + VP2C-GFP-mRuby3) |
| yEGFP_VP2_fw | yeGFP forward, with overhang for VP2C linker | GGCTCCGGTGGCGGCGGTTCTATGGGATCCTCTAAAGGTGAAG | 43 | pILGFPB5A (Peng et al. 2015[9]) | GFP for VP2C | Linked (ΔVP1 + VP2C-GFP-mRuby3); Coexpressed (ΔVP1 + VP2C-GFP + VP2C-mRuby3) |
| yEGFP_AG4TGGA_rv | yeGFP reverse for GFP-mRuby3 fusion protein; appends a linker peptide (RSAGGGGTGGAEL) sequence | AGCACCACCAGTACCTCCTCCACCAGCAGATCTTTTGTACAATTCATCCATACCATG | 57 | pILGFPB5A | GFP for Linked | Linked (ΔVP1 + VP2C-GFP-mRuby3) |
| yEGFP_GAL10t_rv | yeGFP reverse, with overhang for GAL10 terminator | GATAGTAAGCTGGCAAATTAAGATCTTTTGTACAATTCATCC | 42 | pILGFPB5A | GFP for Coexpressed | Coexpressed (ΔVP1 + VP2C-GFP + VP2C-mRuby3) |
| GAL10t_yEGFP_fw | Forward primer for GAL10 terminator-GAL7 promoter region (for Coexpressed) | CAAAAGATCTTAATTTGCCAGCTTACTATCCTTC | 34 | Yeast strain S288c genomic DNA | GAL10 terminator-GAL7 promoter region | Coexpressed (ΔVP1 + VP2C-GFP + VP2C-mRuby3) |
| GAL7p_VP2_rv | Reverse primer for GAL10 terminator-GAL7 promoter region (for Coexpressed) | GACCACCATTACCCATTTTATCGATTTTTGAGGGAATATTCAACTGTT | 48 | Yeast strain S288c genomic DNA | GAL10 terminator-GAL7 promoter region | Coexpressed (ΔVP1 + VP2C-GFP + VP2C-mRuby3) |

| | | | | | | |
|---|---|---|---|---|---|---|
| ERG20_VP2_fw | FPPS forward, with overhang for VP2C linker. Same primer used for replacing wtFPPS with GPP-overproducing FPPS (F96W-N127W mutant). | CGGTGGCGGCGGTTCTGGATCCATGGCTTCAGAAAAAGAAATTAGG | 46 | pJT9R (Peng et al. 2017[10]) for wtFPPS / pIT6EG7M (Peng et al. 2022[11]) for FPPS(F96W-N127W) | FPPS for VP2C | Linked VLP, Coexpressed VLP |
| ERG20_AG4TGGA_rv | FPPS reverse for FPPS-NES fusion protein; appends a linker peptide (RSAGGGGTGGAEL) sequence. Same primer used for replacing wtFPPS with GPP-overproducing FPPS. | AGCACCACCAGTACCTCCTCCACCAGCAGATCTTTTGCTTCTCTTGTAAACTTTGTTC | 58 | pJT9R for wtFPPS / pIT6EG7M for FPPS(F96W-N127W) | FPPS for Linked | Linked VLP, Linked Free |
| AcNES1_AG4TGGA_fw | NES forward for FPPS-NES fusion protein; appends a linker peptide (RSAGGGGTGGAEL) sequence | AGGAGGTACTGGTGGTGCTGAGCTCATGGCTACCGCAGCAGGT | 43 | pJT9R | NES for Linked | Linked VLP, Linked Free |
| AcNES1_HIS5t_rv | NES reverse, with overhang for HIS5 terminator | AACTGTACATATACTGTTTAAATTAATCTATCTCGAGTTATAAAGATGTGTTATAGATCA | 60 | pJT9R | NES | Linked VLP, Linked Free, Coexpressed VLP, Coexpressed Free |
| ERG20_tGAL10_rv | FPPS reverse for Coexpressed Free, with overhang for GAL10 terminator. Same primer used for replacing wtFPPS with GPP-overproducing FPPS. | CAAGAAGGATAGTAAGCTGGCAAAAGATCTTTATTTGCTTCTCTTGTAAACTTTGTTC | 58 | pJT9R for wtFPPS / pIT6EG7M for FPPS(F96W-N127W) | FPPS | Coexpressed VLP, Coexpressed Free |
| tGAL10_ERG20_fw | Forward primer for GAL10 terminator-GAL7 promoter region (for Coexpressed), with overhang for FPPS | GAACAAAGTTTACAAGAGAAGCAAATAAAGATCTTTTGCCAGCTTACTATCCTTCTTG | 58 | Coexpressed (GFP + mRuby3) | GAL10 terminator-GAL7 promoter region | Coexpressed VLP, Coexpressed Free |
| VP2_AcNES1_rv | VP2C forward (for Coexpressed VLP), with overhang for NES | CCTGCTGCGGTAGCCATGAGCTCAGAACCGCCGCCACCG | 39 | Coexpressed (GFP + mRuby3) | GAL10 terminator-GAL7 promoter-VP2C region | Coexpressed VLP |
| AcNES1_VP2_fw | NES forward (for Coexpressed VLP), with overhang for VP2C | CGGTGGCGGCGGTTCTGAGCTCATGGCTACCGCAGCAGGT | 40 | pJT9R | NES | Coexpressed VLP |

| | | | | | | |
|---|---|---|---|---|---|---|
| ERG20_pGAL10_fw | FPPS forward (for Coexpressed Free), with overhang for GAL10 promoter. Same primer used for replacing wtFPPS with GPP-overproducing FPPS. | AAAGTAAGAATTTTTGAAAATTCAATATAAGGATCCAAAATGGCTTCAGAAAAAGAAATT | 60 | pJT9R for wtFPPS / pIT6EG7M for FPPS(F96W-N127W) | FPPS | Coexpressed Free |
| AcNES1_pGAL7_fw | NES forward (for Coexpressed Free), with overhang for GAL7 promoter | GATAAAAAAAAACAGTTGAATATTCCCTCAAAAGAGCTCAAAATGGCTACCGCAGCAGGT | 60 | pJT9R | NES | Coexpressed Free |
| pGAL7_AcNES1_rv | Reverse primer for GAL10 terminator-GAL7 promoter region (for Coexpressed Free), with overhang for NES | CTGCTGCGGTAGCCATTTTGAGCTCTTTTGAGGGAATATTCAACTGTTTT | 50 | Coexpressed (GFP + mRuby3) | GAL10 terminator-GAL7 promoter region | Coexpressed Free |
| Fseq_GAL10p | Forward sequencing primer for VP2C-cargo; binds to GAL10 promoter | GTGGTAATGCCATGTAATATGATTATTAAAC | 31 | NA | NA | All constructs |
| Rseq_KlURA3t | Reverse sequencing primer for VP2C-cargo; binds to *K. lactis* URA3 terminator | GCATTGGCACGGTGCAACAC | 20 | NA | NA | All constructs |
| Fseq_GAL7p | Forward sequencing primer for Coexpressed; binds to GAL7 promoter | TAGTATTCGTTTGGTAAAGTAGAGG | 25 | NA | NA | Coexpressed VLP, Coexpressed Free |
| Rseq_GAL10t | Reverse sequencing primer for Coexpressed; binds to GAL10 terminator | CTCAACAGTGCTCCGAAG | 18 | NA | NA | Coexpressed VLP, Coexpressed Free |

**Table S3. Synthetic gene sequences.**

| Name | Description | Sequence (5' - 3') | Length (nt) |
|---|---|---|---|
| ΔVP1 (or deltaVP1) | Murine polyomavirus VP1 with the nuclear-localisation signal deleted. Codon-optimised for *S. cerevisiae*. From Cheah et al. 2021[3]. | ATGGCTTCAGGTGTAAGTAAATGCGAAACCAAATGCACAAAGGCTTGCCCTAGACCAGCTCCAGTTCCTAAGTT<br>ATTGATTAAAGGTGGTATGGAAGTTTTGGATTTGGTTACAGGTCCAGATTCTGTTACTGAAATCGAAGCATTTTT<br>GAACCCAAGAATGGGTCAACCACCAACACCAGAATCTTTAACTGAAGGTGGTCAATATTACGGTTGGTCAAGAG<br>GTATTAATTTGGCTACATCTGATACTGAAGATTCACCAGGTAATAATACATTACCAACTTGGTCTATGGCAAAAT<br>TGCAATTACCAATGTTGAACGAAGATTTGACTTGTGATACTTTGCAAATGTGGGAAGCAGTTTCAGTTAAAACAG<br>AAGTTGTTGGTTCTGGTTCTTTGTTGGATGTTCATGGTTTTAATAAGCCAACAGATACTGTTAACACAAAGGGTAT<br>TTCTACTCCAGTTGAAGGTTCACAATATCATGTTTTTGCTGTTGGTGGTGAACCATTGGATTTGCAAGGTTTAGTT<br>ACAGATGCAAGAACTAAGTACAAGGAAGAAGGTGTTGTTACTATTAAAACAATCACTAAGAAAGATATGGTTAA<br>TAAGGATCAAGTTTTGAACCCAATTTCAAAGGCTAAATTGGATAAGGATGGCATGTACCCAGTTGAAATTTGGC<br>ATCCAGATCCAGCTAAAAATGAAAACACAAGATACTTCGGTAATTACACTGGTGGTACTACAACTCCACCAGTT<br>TTGCAATTCACTAACACTTTGACAACTGTTTTGTTAGATGAAAATGGTGTTGGTCCATTATGTAAAGGTGAAGGT<br>TTGTACTTATCTTGCGTTGATATTATGGGTTGGAGAGTTACAAGAAACTACGATGTTCATCATTGGAGAGGTTTG<br>CCAAGATACTTCAAGATTACTTTAAGAAAAAGATGGGTTAAAAATCCATACCCAATGGCTTCTTTGATTTCTTCT<br>TTGTTTAATAATATGTTACCACAAGTTCAAGGTCAACCAATGGAAGGTGAAAACACACAAGTTGAAGAAGTTAG<br>AGTTTACGATGGTACTGAACCAGTTCCAGGTGACCCTGACATGACAAGATATGTAGATAGATTCGGTAAAACTA<br>AAACTGTATTCCCTGGTAATTAA | 1143 |
| VP2C-$[GGGGS]_3$ | Murine polyomavirus VP2(251-301) fused to a $[GGGGS]_3$ linker. Codon-optimised for *S. cerevisiae*. From Cheah et al. 2021[3]. | ATGGGTAATGGTGGTCCAACTCCAGCTGCTCATATTCAAGATGAATCTGGTGAAGTTATTAAATTTTATCAAGCT<br>CAAGTTGTTTCTCATCAAAGAGTTACTCCAGATTGGATGTTGCCATTGATTTTGGGTTTGTATGGTGACATCACTC<br>CAGGTGGTGGTGGTTCTGGTGGCGGTGGCTCCGGTGGCGGCGGTTCT | 198 |
| mRuby3 | Red fluorescent protein mRuby3. Codon-optimised for *E. coli*. From Dashti et al. 2018[12]. | ATGGTGTCTAAGGGCGAAGAGCTGATCAAGGAAAATATGCGTATGAAGGTGGTCATGGAAGGTTCGGTCAACGG<br>CCACCAATTCAAATGCACAGGTGAAGGAGAAGGCAGACCGTACGAGGGAGTGCAAACCATGAGGATCAAAGTC<br>ATCGAGGGAGGACCCCTGCCATTTGCCTTTGACATTCTTGCCACGTCGTTCATGTATGGCAGCCGTACTTTTATCA<br>AGTACCCGGCCGACATCCCTGATTTCTTTAAACAGTCCTTTCCTGAGGGTTTTACTTGGGAAAGAGTTACGAGAT<br>ACGAAGATGGTGGAGTCGTCACCGTCACGCAGGACACCAGCCTTGAGGATGGCGAGCTCGTCTACAACGTCAAG<br>GTCAGAGGGGTAAACTTTCCCTCCAATGGTCCCGTGATGCAGAAGAAGACCAAGGGTTGGGAGCCTAATACAGA<br>GATGATGTATCCAGCAGATGGTGGTCTGAGAGGATACACTGACATCGCACTGAAAGTTGATGGTGGTGGCCATC<br>TGCACTGCAACTTCGTGACAACTTACAGGTCAAAAAAGACCGTCGGGAACATCAAGATGCCCGGTGTCCATGCC<br>GTTGATCACCGCCTGGAAAGGATCGAGGAGAGTGACAATGAAACCTACGTAGTGCAACGCGAAGTGGCAGTTG<br>CCAAATACAGCAACCTTGGTGGTGGCATGGACGAGCTGTACAAGTAA | 714 |

| | | | |
|---|---|---|---|
| NES (or AcNES1) | *Actinidia chinensis* linalool / nerolidol synthase. Codon-optimised for *S. cerevisiae*. PCR-amplified from plasmid pJT9R (Peng et al. 2017[10]) | ATGGCTACCGCAGCAGGTCCTATCGCAACTAACAACTCCCCACAAAACTCCAACGCTTACAGAACTCCAATCGC<br>TCCTTCCGTACCAATTACTCATAAATGGTCTATAGCTGAAGATTTGACATGTATTTCCAATCCTAGTAAGCACAAT<br>AACCCTCAAACTGGTTACAGATCATTTTCTGACGAATTATACGTTAAGTACGAAGAAAAGTTGGAAGATGTTAG<br>AAAAGCATTAAGAGAAGTTGAAGAAAACCCTTTGGAAGGTTTAGTTATGATAGACGCTTTGCAAAGATTGGGTA<br>TCGATTACCATTTCAGAGGTGAAATTGGTGCATTCTTGCAAAAGCAACAAATCATATCTTCAACTCCAGATGGTT<br>ACCCTGAACATGGTTTGTACGAAGTTTCAACATTGTTTAGATTCTTAAGACAAGAAGGTCACAATGTTACCGCTG<br>ACGTCTTTAATAACTTCAAGGATAAGGAAGGTAGATTCAGATCAGAATTGTCAACAGATATTAGAGGTTTGATGT<br>CCTTATACGAAGCAAGTCAATTGAGAATAGAAGGTGAAGACATCTTAGATCAAGCTGCTGATTTCTCCAGTCAAT<br>TGTTAGGTAGATGGACAAAAGATCCTAATCATCACGAAGCCAGATTGGTTTCTAACACTTTAACACATCCATACC<br>ACAAGTCATTGGCTACCTTTATGGGTCAAAAATTGTCCTACATGAACTGCAAGGGTCCAAACTGGGACGGTGTCG<br>ATAATTTGCAAGAATTAGCTAAGATGGATTTGACTATCGTACAAAGTATCCATCAAAAGGAAGTATTCCAAGTTT<br>CTCAATGGTGGAAGGATACAGGTTTAGCCAATGAATTGAAATTGGCTAGAAACCAACCATTGAAGTGGTATATG<br>TGGCCTATGGCCGCTTTAACCGATCCAAGATTCAGTGAAGAAAGAGTTGAATTAACTAAGCCTATTTCTTTTATA<br>TACATCATAGATGACATCTTCGACGTCTATGGTACCATTGAAGAATTGACCTTGTTTACTGATGCAGTTAATAGA<br>TGGGAATTGTCTGCCGTCGAACAATTACCAGACTACATGAAAGTATGTTTCAAGGCTTTGTACGATGTTACCAAC<br>GAAATCGCATACAAAATCTATAAAAAGCATGGTCAAAACCCTATTGATTCCTTGCAAAAGACTTGGGCTAGTTTG<br>TGCAATGCATTTTTAGTTGAAGCAAAGTGGTTCGCCTCTGGTCACTTGCCAAATGCAGAAGAATACTTAAAGAAC<br>GGTATCATCTCTTCAGGTGTTCATGTTGTCTTGGCCCACATGTTTTTCTTGTTAGGTGACGGTATTACACAAGAAT<br>CAGTTGATTTGGTAGATGACTATCCAGGTATTTCCACAAGTATCGCAACCATTTTGAGATTATCTGATGACTTGG<br>GTTCAGCCAAAGATGAAGACCAAGATGGTTATGATGGTTCTTACATCGAATGTTACATGAAGGAACATAAGGGT<br>TCCAGTGTCGATTCAGCCAGAGAAGAAGTAATAAGAATGATCTCCGAAGCATGGAAATGTTTGAATAAGGAATG<br>CTTATCACCAAACCCTTTTTCTGAATCATTCAGAATAGGTTCCTTGAATATGGCTAGAATGATCCCTATGATGTAC<br>TCTTACGATGACAACCATAACTTGCCAATTTTAGAAGAACACATGAAGGCAATGATCTATAACACATCTTTATAA | 1722 |
| FPPS (F96W-N127W) | *S. cerevisiae* farnesyl diphosphate synthase (ERG20) with the F96W-N127W double point mutations.<br><br>PCR-amplified from plasmid pIT6EG7M (Peng et al. 2022[11]). | ATGGCTTCAGAAAAAGAAATTAGGAGAGAGAGATTCTTGAACGTTTTCCCTAAATTAGTAGAGGAATTGAACGC<br>ATCGCTTTTGGCTTACGGTATGCCTAAGGAAGCATGTGACTGGTATGCCCACTCATTGAACTACAACACTCCAGG<br>CGGTAAGCTAAATAGAGGTTTGTCCGTTGTGGACACGTATGCTATTCTCTCCAACAAGACCGTTGAACAATTGGG<br>GCAAGAAGAATACGAAAAGGTTGCCATTCTAGGTTGGTGCATTGAGTTGTTGCAGGCTTACTGGTTGGTCGCCGA<br>TGATATGATGGACAAGTCCATTACCAGAAGAGGCCAACCATGTTGGTACAAGGTTCCTGAAGTTGGGGAAATTG<br>CCATCTGGGACGCATTCATGTTAGAGGCTGCTATCTACAAGCTTTTGAAATCTCACTTCAGAAACGAAAAATACT<br>ACATAGATATCACCGAATTGTTCCATGAGGTCACCTTCCAAACCGAATTGGGCCAATTGATGGACTTAATCACTG<br>CACCTGAAGACAAAGTCGACTTGAGTAAGTTCTCCCTAAAGAAGCACTCCTTCATAGTTACTTTCAAGACTGCTT<br>ACTATTCTTTCTACTTGCCTGTCGCATTGGCCATGTACGTTGCCGGTATCACGGATGAAAAGGATTTGAAACAAG<br>CCAGAGATGTCTTGATTCCATTGGGTGAATACTTCCAAATTCAAGATGACTACTTAGACTGCTTCGGTACCCCAG<br>AACAGATCGGTAAGATCGGTACAGATATCCAAGATAACAAATGTTCTTGGGTAATCAACAAGGCATTAGAACTT<br>GCTTCCGCAGAACAAAGAAAGACTTTAGACGAAAATTACGGTAAGAAGGACTCAGTCGCAGAAGCCAAATGCA<br>AAAAGATTTTCAATGACTTGAAAATTGAACAGCTATACCACGAATATGAAGAGTCTATTGCCAAGGATTTGAAG<br>GCCAAAATTTCTCAGGTCGATGAGTCTCGTGGCTTCAAAGCTGATGTCTTAACTGCGTTCTTGAACAAAGTTTAC<br>AAGAGAAGCAAA | 1056 |
